# *Aspergillus* dsRNA virus drives fungal fitness and pathogenicity in the mammalian host

**DOI:** 10.1101/2024.02.16.580674

**Authors:** Marina Campos Rocha, Vanda Lerer, John Adeoye, Hilla Hayby, Maria Laura Fabre, Amelia E. Barber, Neta Shlezinger

**Affiliations:** Koret School of Veterinary Medicine, Faculty of Agriculture, The Hebrew University, Rehovot, Israel; Institute of Microbiology, Friedrich Schiller University, Jena, Germany

**Keywords:** mycovirus, hypervirulence, polymycovirus, *Aspergillus fumigatus*, viability, host-pathogen interaction

## Abstract

Fungal pathogens pose a significant threat to global health. *Aspergillus fumigatus* accounts for approximately 65% of all invasive fungal infections in humans, with mortality rates from invasive aspergillosis reaching nearly 50%. Mycoviruses, viruses that infect fungi, can modulate fungal virulence in plant pathogenic fungi, leading to either hypovirulence or hypervirulence. However, their impact on fungal pathogenesis in mammals has remained largely unexplored. Here, utilizing an *A. fumigatus* strain naturally infected with Aspergillus fumigatus polymycovirus*-1M* (AfuPmV-1M), we found that the mycovirus confers a significant survival advantage to the fungus under conditions of oxidative stress, heat stress, and within the murine lung. Thus, AfuPmV-1M modulates fungal fitness, resulting in increased virulence and the progression of exacerbated fungal disease. Moreover, antiviral treatment reverses the virus-mediated increase in virulence, representing a promising “antipathogenicity” therapy against virus-bearing pathogenic fungi. Collectively, these findings reveal that mycoviruses act as pivotal’backseat drivers’ in human fungal diseases, underscoring significant clinical implications and offering promising avenues for novel therapeutic strategies.

## Main Text

Human fungal pathogens contribute to a significant annual global burden, resulting in an estimated 1 billion infections and over 1.6 million deaths. This mortality rate is comparable to tuberculosis and surpasses that of malaria and breast cancer^1^. Mortality rates from invasive fungal infections (IFI) currently stand at approximately 50%, even with standard care. Furthermore, epidemiologists anticipate a consistent rise in IFI cases due to the growing population of individuals with compromised immune systems^2^. Pathogenic filamentous fungi, including *Aspergillus fumigatus*, are commonly found in soil and have a ubiquitous presence. These fungi produce a substantial quantity of conidia, which are then released into the air and dispersed by wind. When inhaled, these conidia have the ability to enter the lower respiratory tract ^3^. In individuals with a competent immune system, the conidia are efficiently eliminated from the lungs. However, immunocompromised patients face a heightened risk of developing acute invasive mycosis. Among all invasive fungal infections in humans, *A. fumigatus* constitutes approximately 65% of the cases, and the mortality rates for disseminated disease can reach up to 50% ^2^.

While fungi play a significant role in causing diseases in humans, animals, and plants, they themselves are not immune to viral infections. Mycoviruses, viruses that infect fungi, are pervasive in the fungal kingdom, infecting over 20% of the tested isolates ^4^.

The overwhelming majority of mycoviruses possess either double-stranded (ds) or single-stranded (ss) RNA genomes, with only one family known to have circular ssDNA genomes ^4^. While certain mycoviral infections exhibit benign characteristics, there are notable instances where mycoviruses have either advantageous or detrimental impacts on their fungal hosts.

These effects encompass a range of outcomes, including modified virulence ^5,6^, drug resistance ^7^, mycotoxin production ^8,9^, and enhanced thermotolerance ^10^. However, the impact of mycoviruses on fungal physiology and virulence, along with the underlying mechanisms mediating these effects, remain inadequately understood. This is particularly true for the intricate interplay among mycoviruses infecting human pathogenic fungi, the fungal host itself, and the mammalian host.

The Af293 strain of *A. fumigatus*, isolated from the lung of a patient who succumbed to aspergillosis, naturally harbors Aspergillus fumigatus polymycovirus-1 (AfuPmV-1). This segmented double-stranded RNA (dsRNA) virus, belonging to the Polymycoviridae family, encodes four open reading frames (ORFs)^11^. Subsequent research identified a closely related variant, AfuPmV-1M, that is distinguished by a fifth ORF of unknown function^12^. Multiple studies have explored the physiological impacts of these mycoviruses on *A. fumigatus*, reporting diverse and often contradictory phenotypes, including altered pigmentation, conidiation, and resistance to antifungals and stress conditions^7,11,13–16^. However, their roles in fungal virulence and pathogenicity remain poorly understood, with limited and conflicting evidence. One study reported AfuPmV-1-induced hypervirulence in a *Galleria mellonella* infection model^11^, while another observed AfuPmV-1M-associated hypovirulence in an immunosuppressed mouse model of invasive pulmonary aspergillosis (IPA)^12^. Beyond *A. fumigatus*, polymycoviruses in plant-and insect-pathogenic fungi have been linked to enhanced virulence^17,18^, and recent reports suggest mycovirus-mediated modulation of mammalian immune responses by clinical isolates of *Talaromyces* and *Malassezia* ^6,19,20^.

These findings collectively suggest that mycoviruses may act as drivers of fungal virulence, contributing to strain-specific differences in pathogenicity and host adaptation. Yet, the mechanisms driving mycovirus-mediated effects on fungal virulence and disease outcomes in mammalian hosts remain largely unresolved, leaving critical gaps in our understanding of their impact. Here, we investigate how mycoviral infection shapes *A. fumigatus* fitness, virulence, and infection outcomes, providing new insights into the complex interplay between endogenous mycoviruses and fungal pathogenicity.

## Results

### AfuPmV-1M Modulates Conidiation in *A. fumigatus*

To investigate the impact of mycoviral infections on *A. fumigatus* physiology, we first confirmed the presence of a polymycovirus within our Af293 laboratory strain (Fig. 1).

**Figure 1.**
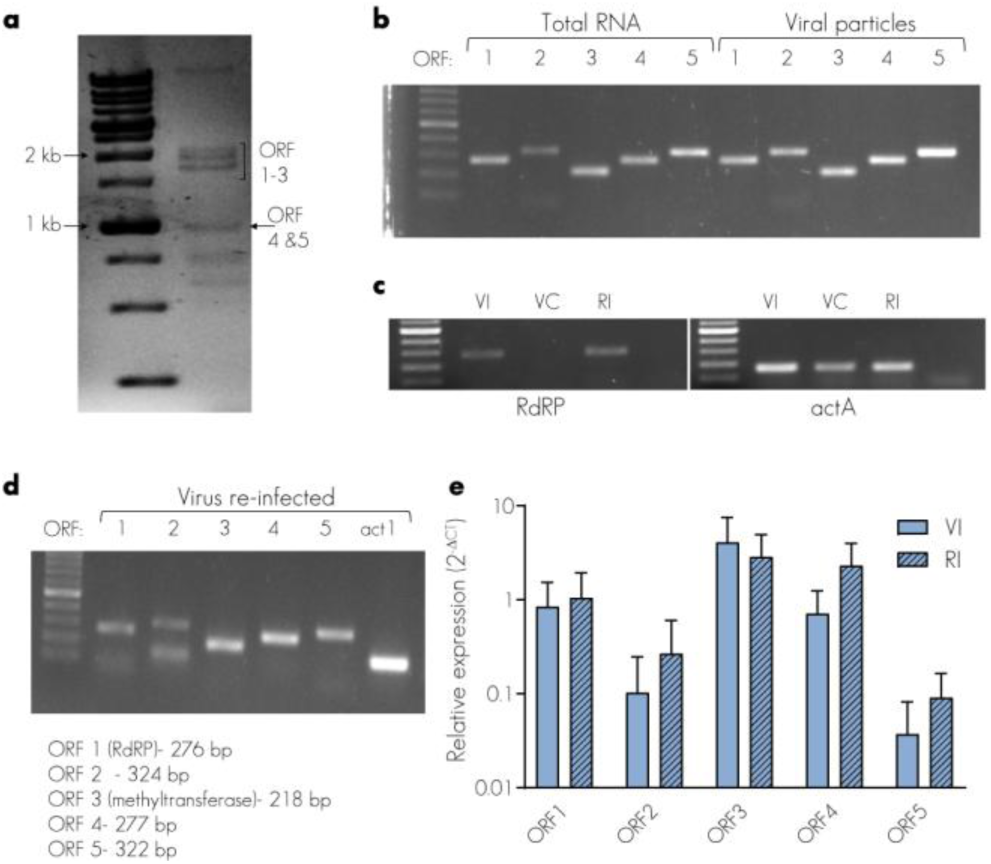
Identification of AfuPmV-1M and generation of congenic virus-cured and re-infected strains. **a.** dsRNA extracted from *A. fumigatus Af293* and treated with DNase I and S1 nuclease was fractionated on a 1% agarose gel. **b.** Reverse-transcription PCR (RT-PCR) amplification to confirm the presence of *AfuPmV-1M* ORF 1-5 in Af293 total RNA and isolated viral particles. **c.** RT-PCR amplification of a 276 bp segment from Afupmv-1M RdRP in total RNA extracted from Af293 (VI), cycloheximide virus-cured (VC), and Re-infected (RI) strains. **d.** RT-PCR amplification of *AfuPmV-1M* ORF 1-5 from total RNA extracted from the re-infected strain. **e.** Real-time qPCR showing relative expression levels of AfuPmV-1M ORF 1-5 (24 h in liquid glucose minimal media). Data are from six biologically independent replicates, normalized to *A. fumigatus actA*. One-way ANOVA with Tukey’s multiple comparison test. Error bars indicate standard error around the mean.

Double-stranded RNA (dsRNA) extracted from Af293 was analyzed using agarose gel electrophoresis, revealing four distinct bands, corresponding to ORF1-4 of AfuPmV-1M (Fig 1a). Notably, ORF 4 and ORF 5 share a similar size of approximately 1150 bp (1140 bp and 1158 bp, respectively), making them indistinguishable using agarose gel alone. To determine whether ORF5 is encoded by the detected virus, we performed reverse transcription PCR (RT-PCR) analysis using both total RNA and RNA from isolated viral particles and primers specific for each ORF (Fig. 1b). This analysis revealed the presence of ORF1-5, confirming AfuPmV-1M as the infecting mycovirus variant in our strain. To generate a congenic virus-cured (VC) strain, we utilized the cycloheximide translation inhibition curing approach^21^.

Subsequently, the virus was reintroduced into the VC strain to generate a re-infected strain (RI) through protoplast transfection with AfuPmV-1M dsRNA ^11^. The purity of the decontaminated strain (VC) and successful reintroduction of ORF1-5 were verified after four successive passages on non-selective GMM agar through reverse transcription PCR and quantitative real-time PCR analysis of total RNA (Fig. 1c-d). Real-time qPCR analysis confirmed that the expression levels of AfuPmV-1M ORFs 1-5 remain stable and comparable in the re-infected strain, relative to the ancestral Af293 strain (Fig. 1e). To confirm that the observed phenotypes are AfuPmV-1M-specific and that the cycloheximide treatment did not lead to detrimental genetic changes, we generated an additional cured strain using the antiviral nucleoside analog ribavirin, as previously described^22^ (Supplementary Fig. 1a).

Additionally, we performed Illumina whole genome sequencing of the cycloheximide-cured, ribavirin-cured, and re-infected strains to screen for genetic changes compared to our lab strain of Af293. No nucleotide changes were identified in any of the derivative strains, and no changes in aneuploidy (i.e. whole chromosome) or copy number variation of smaller regions were detected in any strain (Supplementary Fig. 1b).

In agreement with previous studies^11,15^, the three congenic strains displayed comparable radial growth, biomass production, biofilm formation, and germination rates in basal conditions (GMM, 37°C) (Supplementary Fig. 2). However, both the original virus-infected (VI) and re-infected (RI) strains exhibited colonies with a darker color, suggesting a decrease in conidiation and/or melanin production in the virus-cured (VC) strains (Fig. 2a, Supplementary Fig. 3a).

**Figure 2.**
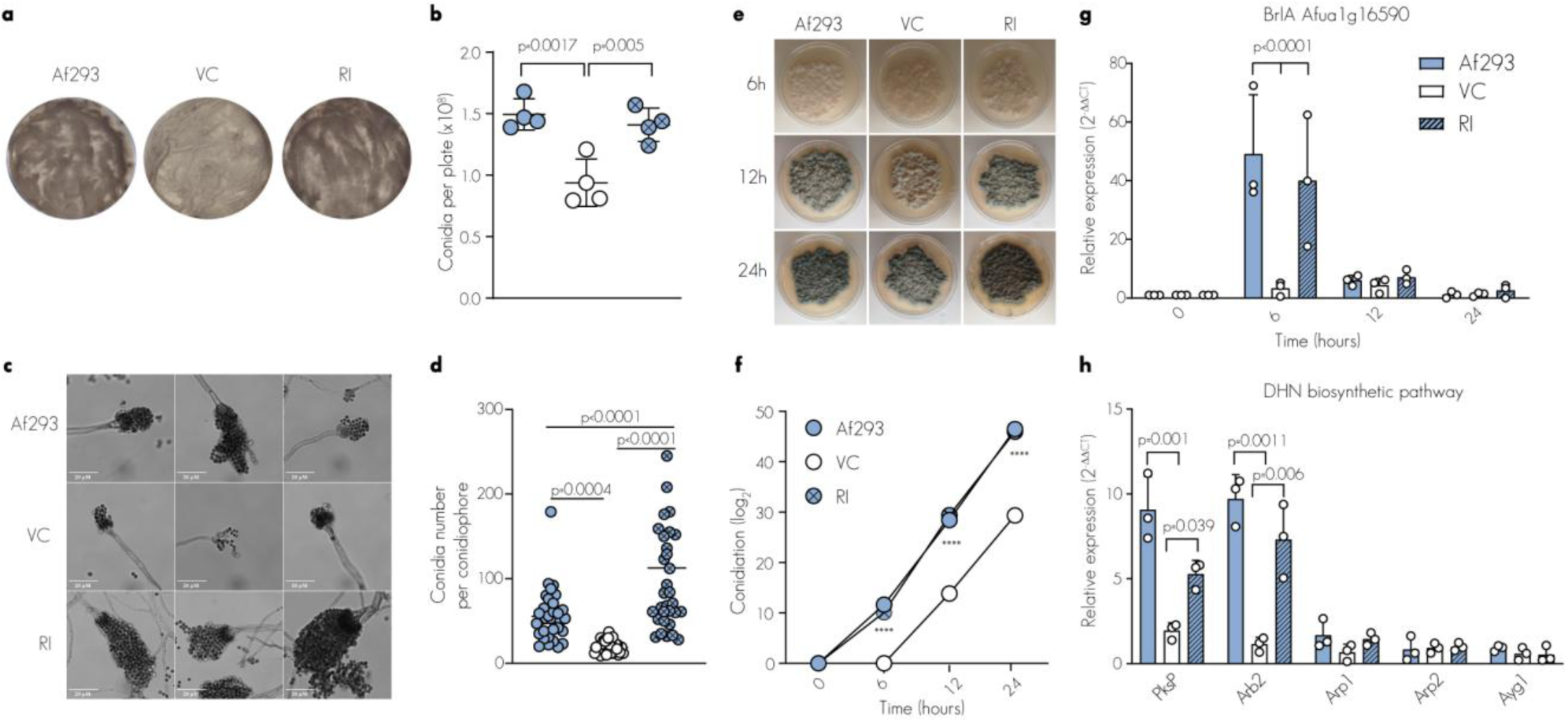
AfuPmV-1M regulation of conidiation and melanin production. **a.** AfuPmV-1M-infected strains (Af293 and RI) exhibit darker colonies compared to the cycloheximide virus-cured strain. 2×10^5^ conidia (CFUs) were spread over GMM plates and grown for 7 days. **b.** Quantification of conidia number per plate on day 7. **c.** Representative micrographs of conidiophores. Images representative of three biologically independent experiments. **d.** Conidia number per conidiophore. **e.** Representative micrographs of synchronized asexual development induction in *A. fumigatus* Af293, VC, and RI. The time (in h) indicates the growth following the transfer of the mycelium from the liquid-submerged synchronized culture to solid medium. **f.** Conidia quantification in a time course experiment of synchronized asexual differentiation. **** <0.0001.**g-h.** Real-time qPCR showing relative expression levels of *brlA* (**g**) and *A. fumigatus* gene cluster involved in DHN-melanin biosynthesis (**h**) during synchronized asexual developmental induction for *A. fumigatus Af293* and congenic strains. RNA was extracted at the indicated time points (**g**) or 12 h (**h**) following the transfer of mycelium from the liquid-submerged synchronized culture to a solid medium. Data were analyzed using the 2^−ΔΔCt^ method. Vegetative mycelia (hyphal state) served as control sample and normalized against 18S expression (reference gene). The data are derived from four (**b**) or three (**c-h**) biologically independent experiments conducted in triplicate. Statistical analysis: (**b,d**) one-way ANOVA and (**f-h**) two-way ANOVA with Tukey’s multiple comparison test. Conidial suspensions were adjusted based on viability staining to account for differences in conidial viability prior to each experiment. Error bars indicate standard error around the mean.

The quantitative analysis of conidia counts per plate and per conidiophore revealed a significant decrease in asexual conidia production per plate, along with a marked reduction in the number of conidia per conidiophore in the virus-cured strain (Fig. 2b-d, Supplementary Fig. 3b). We then further investigated the conidiation defect in the virus-cured strain during synchronized asexual differentiation. Conidia production was monitored up to 24 h after the initiation of asexual conidiation. The results showed that the virus-infected strains (Af293 and RI) produced conidia by 6 h, whereas conidiation in the virus-cured strain was delayed, initiating between 6 and 12 h after induction. This represents a significant delay in conidia formation in the mycelia of the virus-cured strain compared to the virus-infected strains (Fig. 2e-f). The conidiation process in *A. fumigatus* is regulated by the central regulatory pathway (CRP), which consists of three essential genes: *brlA*, *abaA*, and *wetA*^23,24^. To determine whether AfuPmV-1M infection is involved in conidiation regulation, we examined the mRNA levels of *brlA*, *abaA*, and *wetA* in both virus-infected (Af293 and RI) and virus-cured strains during conidiogenesis (Fig. 2g, Supplementary Fig. 3c-d). We observed a significant reduction in *brlA* expression in the virus-cured strain during the early stages of conidiation, 6 h after the induction of conidiation. These results suggest that AfuPmV-1M is essential for the proper regulation of conidiation in the Af293 strain background. Interestingly, in addition to its role in conidiation, BrlA serves as a master regulator of genes involved in the biosynthesis of secondary metabolites, including gliotoxin and DHN-melanin, both of which contribute to the pathogenicity of this human pathogen^25,26^. Indeed, in addition to the altered conidiation and consistent with the lighter appearance on plates (Fig. 2a), the expression of the polyketide synthase gene *pksP*, a key enzyme in DHN-melanin biosynthesis, and *Arb2*, a key regulator of melanogenesis, was 5 to 8 times higher (for *pksP*) and 6 to 10 times higher (for *Arb2*) in the virus-infected strains compared to the virus-cured strains, 12 h after conidiation induction (Fig. 2h, Supplementary Fig. 3e-i). This establishes a clear relationship between the expression of DHN-melanin biosynthesis genes and the presence of AfuPmV-1M.

### AfuPmV-1M Enhances Survival During Sporulation and Environmental Stresses

As both a compost-dwelling organism and an opportunistic human pathogen, *A. fumigatus* must endure variable and often extreme environmental conditions, including the elevated temperatures (often exceeding 50°C), and fluctuating pH of composting soil and the oxidative, and hostile environment of the phagosome within host immune cells. To assess the impact of AfuPmV-1M on stress tolerance, we compared the susceptibility of the virus-infected and virus-cured strains under various stress conditions (Fig. 3, Supplementary Fig. 4).

**Figure 3.**
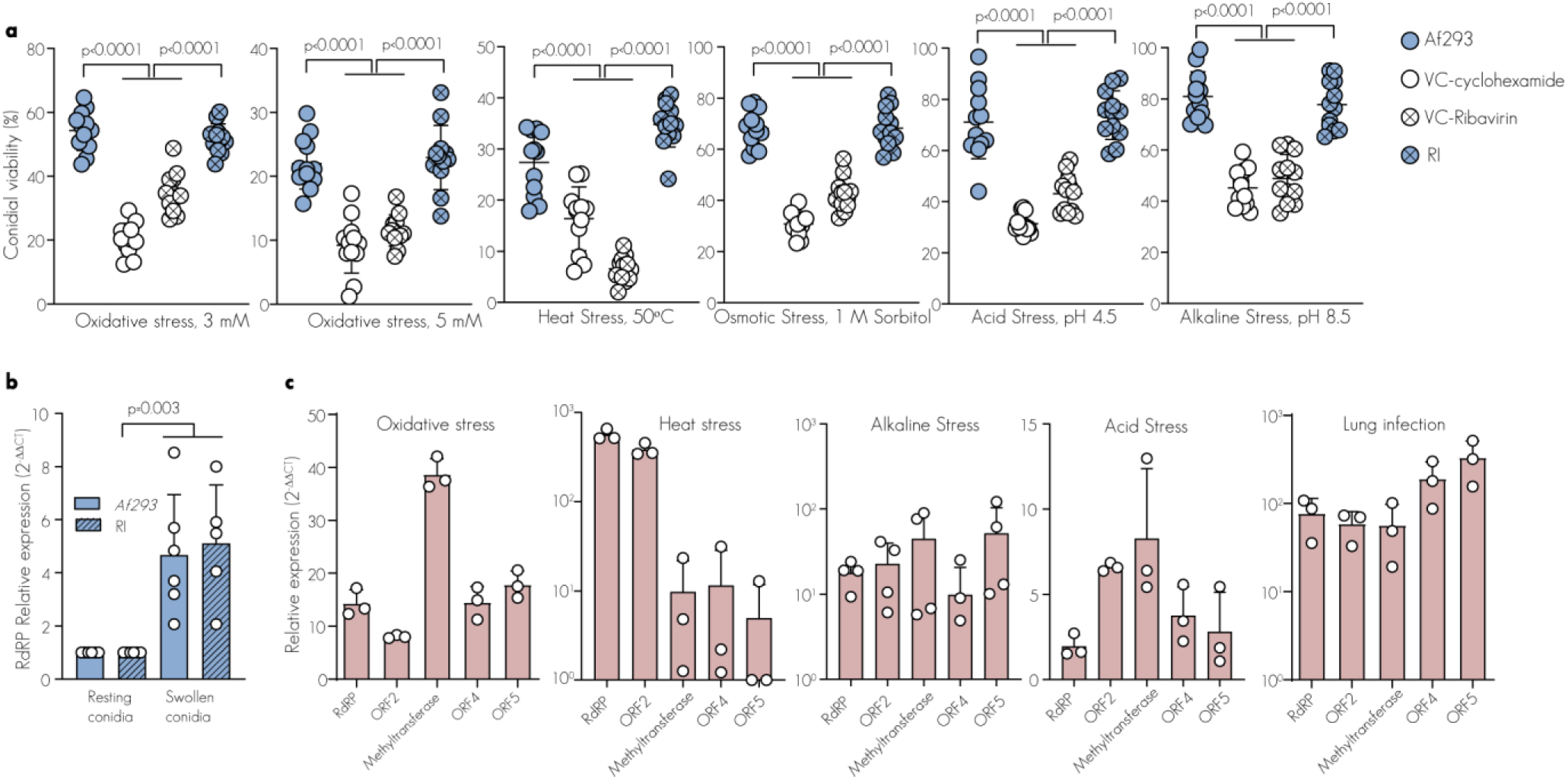
AfuPmV-1M Mediates Cytoprotective Response to Environmentally-and Infection-Relevant Stresses. **a.** Conidial viability under stress conditions. 2×10^5^ *Af293*, cycloheximide virus-cured, ribavirin virus-cued, and virus-re-infected (RI) conidia (CFUs) swollen conidia (4h) were exposed to 3 or 5 mM H_2_O_2,_ 50°C, 1 M sorbitol, pH 4.5 or pH 8.5 for additional 4 h and analyzed for CFUs. **b.** Real-time qPCR showing relative expression levels of AfuPmV-1M RdRP (ORF1) in resting conidia harvested from 4-day old plates and swollen conidia (4 h at 37°C). Data were analyzed using the 2^−ΔΔCt^ method with Af293 or RI resting conidia as the control sample for the corresponding swollen conidia sample and normalized against *ActA* expression (reference gene). **c.** Real-time qPCR showing relative expression levels of AfuPmV-1M ORF 1-5 upon oxidative stress (5 mM, 4h), heat shock (50°C, 4 h), alkaline stress (pH 8.5, 4 h), acid stress (pH 4.5 4 h) and during infection in IPA infection model (24 h p.i.). Data were analyzed using the 2^−ΔΔCt^ method with physiological conditions (GMM broth, 37°C, 24 h) as the control sample and normalized against *ActA* expression (reference gene). The data are derived from twelve (**a**), six (**b**) or three (**c**) biologically independent experiments. Statistical analysis: (**a**-**b**) one-way ANOVA with Tukey’s multiple comparison test. Conidial suspensions were adjusted based on viability staining to account for differences in conidial viability prior to each experiment. Error bars indicate standard error around the mean.

The three strains exhibited similar growth and survival under basal conditions and in the presence of the ergosterol biosynthesis–targeting antifungal drug voriconazole (Supplementary Fig. 2 and 4). However, conidia from the mycovirus-carrying strains demonstrated heightened resistance to oxidative challenge representing the phagosomal environment ^27^, as well as to elevated temperatures, low and high pH, and osmotic challenge—mimicking the natural habitat of composting soil ^28,29^ (Fig. 3a, Supplementary Fig. 4). These data indicate that in the absence of the virus, the fungus is less competitive in its natural habitat, such as compost piles where *A. fumigatus* is commonly found at temperatures exceeding 50°C ^30^, as well as in the lung environment where fungal cells are exposed to oxidative stress from host immune cells ^31^. Notably, the cytoprotective effect of AfuPmV-1M was most pronounced when swollen conidia were challenged, although resting virus-infected conidia also showed enhanced survival rates compared to virus-cured resting conidia under oxidative, heat, low and high pH, and osmotic stress conditions, albeit to a lesser extent (Supplementary Fig. 4). Consistent with this observation, quantification of mycoviral RdRP (transcription and genomic dsRNA) in swollen and resting conidia revealed higher RNA levels in the swollen conidia of Af293 and the congenic re-infected strain, supporting the notion that AfuPmV-1M mediates the cytoprotective effect (Fig. 3b).

Remarkably, mycoviral total RNA (transcript and genomic) was upregulated under these conditions as assessed by RT-qPCR of ORF1-5 under oxidative stress and heat shock, low and high pH, and during infection in a murine pulmonary invasive aspergillosis model (Fig. 3c). These results suggest that stress conditions may activate the mycovirus. Given that the virus-infected strains displayed enhanced resistance to these stressors, it is plausible that mycoviral activation modulates fungal pathways that promote the survival of its fungal host as a mechanism of self-preservation.

To elucidate the underlying mechanism through which AfuPmV-1M influences fungal fitness, we conducted a comprehensive investigation of mycovirus-derived proteomic alterations under both basal conditions and oxidative stress (H_2_O_2_ 8 mM), employing proteomic profiling (Fig. 4, Supplementary Fig. 5, and Dataset 1). In this analysis, we quantified abundances for between 3,390 and 3,525 proteins. Applying filters of a p-value < 0.05 and a fold change log2 cutoff of ≥0.9 or ≤-0.9 revealed that under basal conditions, the AfuPmV-1M-infected strain Af293 displayed significant alterations in protein levels compared to the congenic virus-cured strain. Specifically, under control conditions, we observed that 17 proteins were increased and 16 were decreased in the AfuPmV-1M-infected strain (Fig. 4a). The most upregulated protein in the presence of AfuPmV-1M is Brf1 (AFUA_3G12730,10-fold), a subunit of the transcription factor TFIIIB complex required for accurate initiation of transcription by RNA Polymerase III (Pol III), which synthesizes transfer RNAs (tRNAs), 5S ribosomal RNA (rRNA), and other essential RNA molecules ^32^. To investigate this, we used RT-qPCR to first determine whether elevated Brf1 protein levels result from increased Brf1 transcript abundance and then assess if this enhances transcription of its non-coding RNAs targets, as expected from Brf1’s role in TFIIIB (Fig. 4b-c). The Pol III genes analyzed included U6 snRNA (U6), tRNA-Arg (Arg), tRNA-Phe (Phe), and tRNA-Tyr (Tyr). Consistent with the proteome data, we observed that under basal conditions, Brf1 and its targets, tRNA-Arg and tRNA-Phe, were higher in both Af293 and re-infected strains compared to the virus-cured strain. However, tRNA-Tyr expression was unchanged, and U6 snRNA was upregulated specifically in the virus-cured strain despite Brf1 upregulation. This suggests additional regulatory factors might influence gene transcription, and different tRNA genes could vary in sensitivity to Brf1 levels^33,34^. The increase in U6 snRNA in the virus-cured strain may indicate a feedback loop modulating its expression^35^. When subjected to oxidative stress, Brf1 expression remained unchanged in the virus-infected strains (Af293 and RI) and was slightly elevated in the virus-cured strain, consistent with the proteome data showing a 12-fold increase in H_2_O_2_-exposed VC compared to VC under physiological conditions. However, its targets—tRNAs Arg, Tyr, and Phe, as well as U6 snRNA—were elevated in the virus-infected strains (Af293 and RI). An additional group of proteins that were enriched in the AfuPmV-1M-infected strain compared to the virus-cured strain are known components of stress granules (SGs). These cytoplasmic condensates rapidly form during various stress conditions, including heat shock, oxidative stress, hypoxia, bacterial, and viral infections ^36–38^. Their dynamic nature is mediated via stored translational preinitiation complexes, which can be rapidly released to resume gene expression when cellular stress subsides. Our proteomic analysis identified known SG components exhibiting increased abundance in an AfuPmV-1M-dependent manner including Ribosomal protein L24 (AFU_5G08770, 3.7-fold), RNase III (AFUA_5G04440, 2.14-fold), polyadenylation factor subunit CstF64 (AFUA_2G09100, 2-fold), Thioredoxin (AFUA_3G14970, 3.33-fold), RNA methyltransferase (AFUA_4G13450, 3.17-fold), and U6 small nuclear ribonucleoprotein (Lsm3) (AFUA_5G12570,2.5-fold, p-value = 0.066066) (Fig. 4a) ^38–45^. In agreement with the proteome data, RT-qPCR analysis demonstrated increased mRNA levels of U6 snRNA, ribosomal protein L24, RNase III, and methyltransferase under controlled basal conditions. Furthermore, the expression of the *Pbp1* gene, which encodes the poly(A) binding protein PAB1—a highly conserved key regulator of mRNA metabolism and a defining marker of stress granules ^46,47^— showed increased abundance in the virus-infected strains (Fig. 4e). To determine whether AfuPmV-1M modulates fungal SGs, we generated congenic virus-infected and virus-cured strains with a chromosomal C-terminal GFP fusion at the endogenous Pab1 locus. Under basal growth conditions (37°C), Pab1-GFP was predominantly cytoplasmic in both virus-infected and virus-cured strains (Fig. 4f). However, the fluorescence intensity of Pab1-GFP was significantly higher in the virus-infected strain, as shown by flow cytometry (Fig. 4g), indicating increased Pab1 expression under physiological conditions. Following a rapid temperature shift to 50°C, Pab1-GFP formed multiple cytoplasmic puncta in both virus-infected and virus-cured strains (Fig. 4f). Notably, the number of Pab1-GFP foci was significantly higher in the virus-infected strain, as quantified using image-based cell clustering analysis via ImageStream (Fig. 4h, Supplementary Fig. 6). The formation of SGs has been demonstrated to play a pivotal role in driving stress adaptation and promoting cell survival^48^. Our findings demonstrate that AfuPmV-1M-infected strains exhibit heightened resistance to oxidative and heat stress. This observation, coupled with the upregulation of SGs in these strains, suggests that AfuPmV-1M promotes SG formation, thereby bolstering the fungal host’s capacity to endure subsequent lethal stress. Moreover, our proteomic analysis revealed that Thioredoxin, a key antioxidant system component in defense against oxidative stress^49^, is significantly elevated at both the transcriptional (38-fold) and translational (4.06-fold) levels in the virus-infected strains compared to the virus-cured strain (Fig. 4a,e). Interestingly, the regulation of the high-affinity iron transporter (FtrA) in *Aspergillus* was also influenced by the presence of the virus. While proteomic data indicated a downregulation of FtrA in the AfuPmV-1M-infected strain, qPCR analysis showed an upregulation at the transcriptional level. This apparent discrepancy might be attributed to post-transcriptional or translational regulation mechanisms that differentially affect FtrA expression at the mRNA and protein levels. These findings suggest that AfuPmV-1M modulates multiple aspects of the fungal host’s stress response and iron homeostasis, potentially enhancing its ability to withstand oxidative stress and adapt to changing environmental conditions.

**Figure 4.**
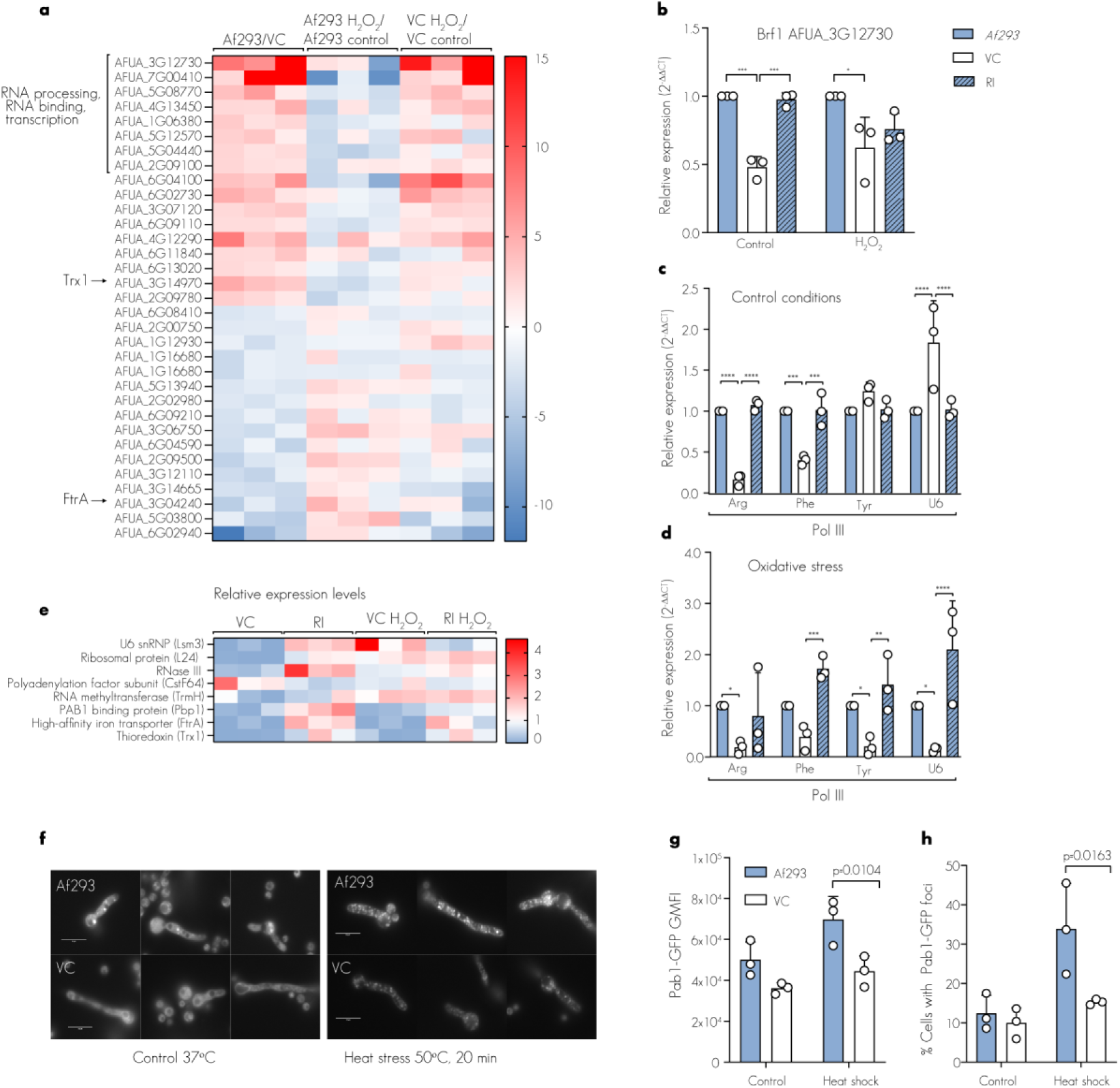
The proteome of AfuPmV-1M-infected *A. fumigatus* is altered. The protein content of *A. fumigatus* Af293 and cycloheximide virus-cured (VC) was characterized using liquid chromatography mass spectrometry (LC-MS/MS) under basal and oxidative stress (8 mM H_2_O_2_, 4h) conditions. **a.** A total of 117 proteins were differentially abundant across the strains and growth conditions, based on an uncorrected value of <0.05 by analysis of variance (ANOVA). n=3. **(b-e)** Relative mRNA levels in Af293 versus VC and RI were analyzed by qRT-PCR using the 2^−ΔΔCt^ method with Af293 as the reference and normalized against 18S rRNA expression (housekeeping gene). **b.** Brf1 expression under control conditions and upon oxidative challenge (8 Mm H_2_O_2_, 4h). (**c-d).** Expression levels of Pol III-transcribed genes: U6 snRNA (U6), tRNA-Arg (Arg), tRNA-Phe (Phe), and tRNA-Tyr (Tyr), under control conditions **(c)** and upon oxidative stress (8 mM H_2_O_2_, 4h) **(d)**. **e**. Relative expression levels of stress-granules components U6 small nuclear ribonucleoprotein (Lsm3) (AFUA_5G12570), Ribosomal protein L24 (AFU_5G08770), RNase III (AFUA_5G04440), polyadenylation factor subunit CstF64 (AFUA_2G09100), RNA methyltransferase (AFUA_4G13450), and PAB1 binding protein (Pbp1) (AFUA_1G09630), High-affinity iron transporter (FtrA) (AFU_5G03800), and Thioredoxin (AFUA_3G14970). n = 3. *<0.04, **< 0.004 ***< 0.0004, **** <0.0001. **f.** Representative fluorescence emission images of congenic PAB1-GFP strains under physiological conditions (37°C) and following heat shock (50°C, 20 min). Conidia were germinated for 7 h before heat shock treatment at 50°C for 20 min **g**. Geometric mean fluorescent intensity (GMFI) of Pab1-GFP expressing strains as analyzed by flow cytometry. **h**. Bar plot showing the percentage of stress-induced Pab1-GFP granule-positive fungal germlings compared to a GFP-expressing control strain. Conidial suspensions were adjusted based on viability staining to account for differences in conidial viability prior to each experiment.

Statistical analysis: (**b-d, g-h**) two-way withTukey’s multiple comparison test. Error bars indicate standard error around the mean (center).

**Table 1-Selected proteins altered in AfuPmV-1M+ proteomes.** Protein content of the AfuPmV-1M-infected and cycloheximide-cured strains was catalogued by mass spectrometry. Data are mean + s.e.m., n=3. The full proteomic analyses can be consulted in supplemental data files (Dataset 1).

### AfuPmV-1M Protects A. fumigatus from Intraphagosomal Killing In vivo Through Intrinsic Cellular Mechanisms

To evaluate the impact of mycoviral infection on fungal fitness in vivo, we conducted a comparative analysis of leukocyte conidial uptake and killing in immunocompetent mice challenged with Af293 (VI), virus-cured (VC), and Re-infected (RI) conidia (Fig. 5a). For this purpose, we employed a mixed infection model in which the three strains were color-coded using an invariant cell wall labeling technique, as previously described ^50,51^. This experimental design allowed us to examine the functional outcomes in the presence of virus-infected and virus-cured conidia within the same inflammatory milieu. By doing so, we could focus specifically on cell-intrinsic-mediated effects, independent of secondary influences on microbial burden and tissue inflammation that often arise when comparing singly-infected mice. At 24 h post-infection, the lungs of infected mice were harvested. Neutrophils containing conidia a single strain were identified and sorted into three distinct neutrophil populations based on their phagocytosis of either VI, VC, or RI conidia (Fig. 5b, Supplementary Fig. 7a). Phagocytosis by neutrophils was not affected by the presence of the mycovirus, suggesting that differences in conidial uptake are unlikely to impact fungal virulence (Supplementary Fig. 7b). However, analysis of colony-forming units (CFUs) in the sorted populations revealed enhanced survival of virus-infected conidia (Af293 and RI) within lung neutrophils (Fig. 5c). Similarly, the virus-infected strains (Af293 and RI) demonstrated heightened resistance to conidial killing by human derived neutrophils, and by the environmental soil-dwelling phagocytes, *Acanthamoeba castellanii*, while phagocytosis remained similar (Fig. 5d-e, Supplementary Fig. 7c). These data provide compelling evidence that AfuPmV-1M bestows protection against neutrophil-mediated conidial killing in the lung. By comparing different strains in the same lung environment, we have established that the cytoprotective effect is cell-intrinsic, ruling out the influence of variations in cell recruitment and inflammatory response.

**Figure 5.**
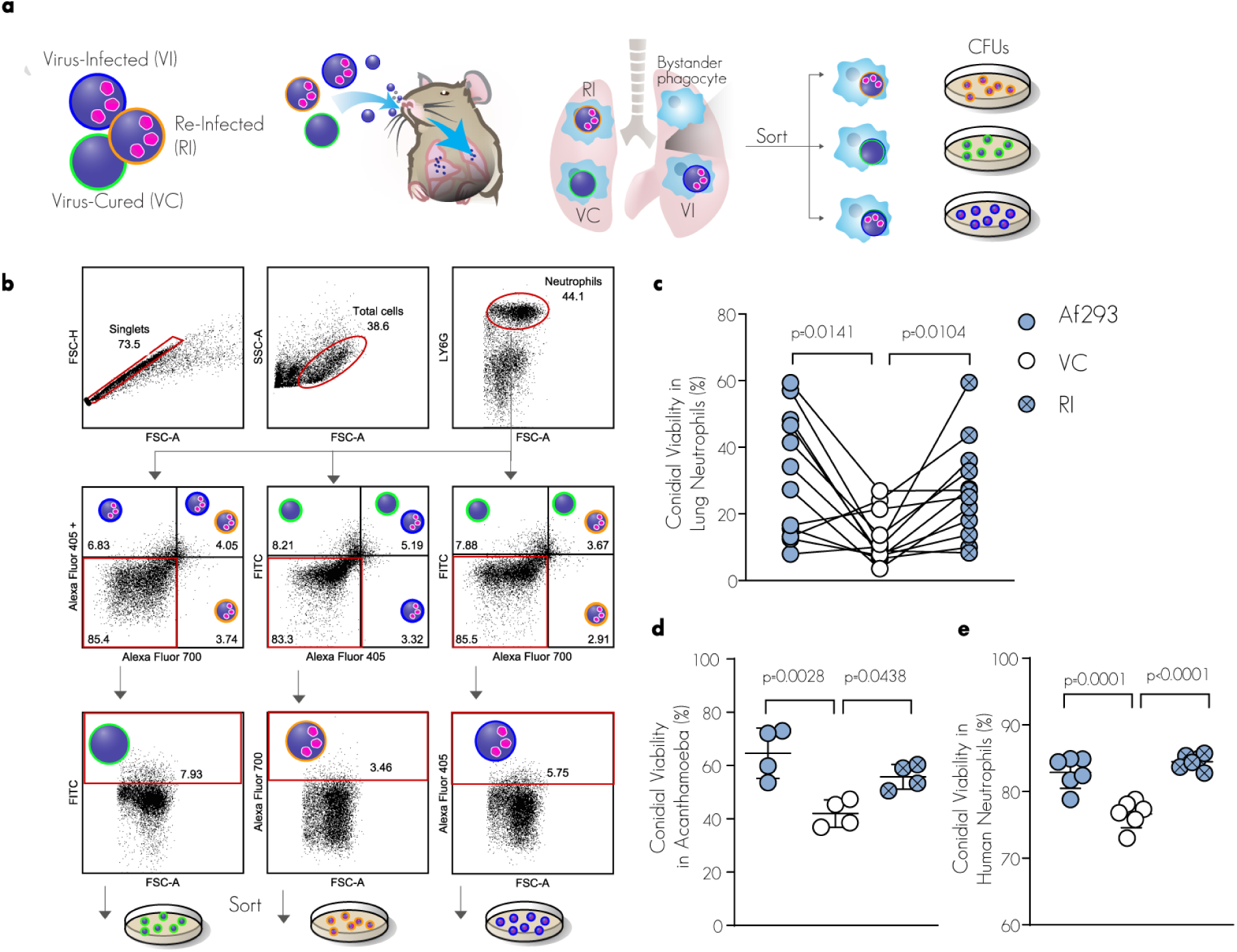
AfuPmV-1M regulates conidial survival in phagocytes. **a.** Mixed infection model. **b.** Gating strategy for sorting neutrophils containing only one conidium of a single strain. Lungs from animals infected with unlabeled conidia were used to define the positive populations. **c.** Fungal viability at 24 h post-infection (p.i.) was calculated based on colony-forming units (CFUs) of sorted neutrophil-phagocytosed strains, including virus-infected, virus-cured (by cycloheximide), and virus re-infected strains. The lines indicate paired data from a single mouse. The data are derived from three biologically independent experiments, each conducted with a cohort of four mice. **d-e.** Swollen conidia were co-incubated with *Acanthamoeba castellanii* **(d)** or human-derived neutrophils **(e)** at MOI of 1:1 for 4 h. Viability was quantified by CFUs. The data are derived from four biologically independent experiments, each performed in triplicate (**d**), or six independent human blood-derived neutrophil samples from two independent experimental repetitions (**e**). Statistical analysis: (**c-e**) one-way ANOVA with Tukey’s multiple comparison test. Conidial suspensions were adjusted based on viability staining to account for differences in conidial viability prior to each experiment. Error bars indicate standard error around the mean (center).

### AfuPmV-1M Promotes Disease Progression

To assess the impact of AfuPmV-1M on virulence, we conducted a challenge experiment using C57BL/6JOlaHsd immunocompetent mice. The mice were inoculated intranasally with 6×10^7^ conidia and the results showed a significantly higher mortality rate in the groups infected with virus-infected strains (Af293 and RI) compared to the group infected with the virus-cured strain (Fig. 6a, Supplementary Fig. 8a). The increased mortality observed in the VI and RI groups correlated with elevated fungal burden in the lungs (Fig. 6b) and tissue damage, as indicated by higher levels of lactate dehydrogenase (LDH) in bronchoalveolar lavage fluid (BALF) at 24 h post-infection (pi) (Fig. 6c). Examination of lung histopathology revealed multiple infection foci and germinating conidia in the virus-infected strain groups (Fig. 6d-e). Following infection with AfuPmV-1M-infected conidia (Af293 and RI), a notable increase in the levels of lung inflammatory cytokines (tumor necrosis factor, IL-1β, IL-6, IFN-β, and IFN-ᵞ) and Myeloperoxidase (MPO) was observed at 72 h post-infection (hpi) in comparison to infection with the congenic virus-cured conidia (Fig. 6f). These findings suggest a higher degree of tissue inflammation and heightened neutrophil activity, respectively. In an ex vivo model using human-derived macrophages challenged with an equal amount of conidia, we did not observe differences in the inflammatory response (Supplementary Fig. 8b). This suggests that the pro-inflammatory response to the virus-infected strain is due to the increased fungal burden. Importantly, the C57BL/6JOlaHsd substrain diverged from the C57BL/6J background approximately 60 years ago and, therefore, presents various phenotypic differences, including susceptibility to infections^52^. Although no previous studies have evaluated the mortality rate of C57BL/6JOlaHsd mice in a pulmonary invasive aspergillosis model, we found that this mouse strain is highly susceptible to *A. fumigatus* Af293 in its competent state, in a mycovirus-dependent manner. To determine whether this mycovirus-mediated virulence is observed in additional mouse models, we challenged ICR and C57BL/6J mice with Af293 and its congenic virus-cured and re-infected strains (Supplementary Fig. 8c-f). We found that although these mice were resistant to infection, the virus-infected strains (Af293 and RI) exhibited higher fungal burden and survived better within the lung environment, while conidial uptake by neutrophils was not affected by the presence of the mycovirus, similar to our observations with C57BL/6JOlaHsd mice. Given that immunocompromised individuals constitute the primary at-risk population for invasive aspergillosis, we challenged cyclophosphamide-treated C57BL/6J mice with 5×10^3^ conidia. We observed significantly higher mortality and fungal burden with the virus-infected strains (Af293 and RI) compared to the virus-cured strain (VC), recapitulating the mycovirus-mediated enhancement of virulence observed in immunocompetent mice (Fig. 6g–h). Taken together, these findings demonstrate that the virus-infected strains exhibit better survival within the lung environment, leading to increased virulence compared to the virus-cured strain, as well as an exacerbated progression of invasive aspergillosis in immunocompetent and immunocompromised mice.

**Figure 6.**
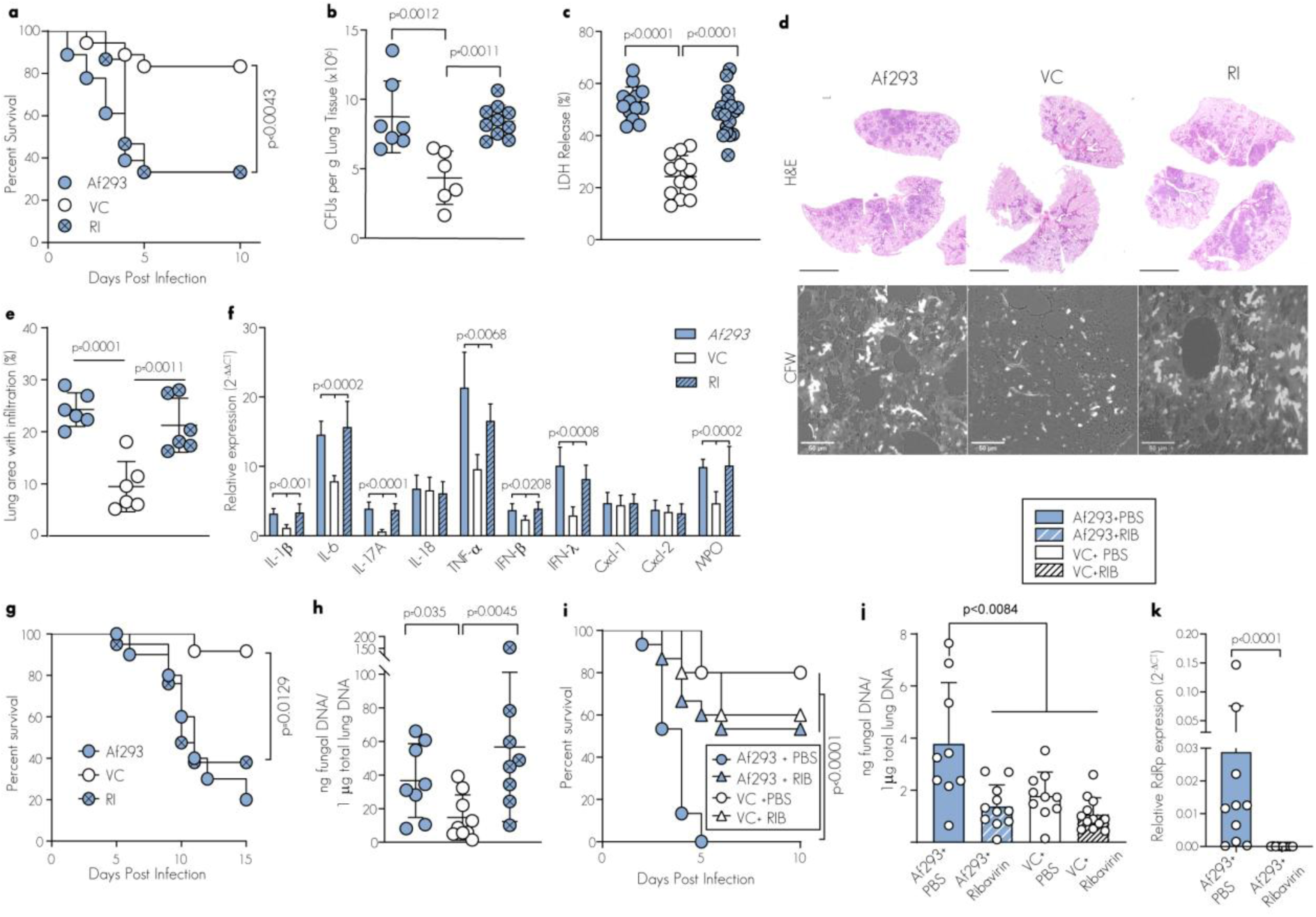
AfuPmV-1 regulates *A. fumigatus* virulence. **a.** Survival of C57BL/6JOlaHsd mice challenged with 6×10^7^ Af293 (n=18), virus-cured by cycloheximide (VC, n=18), or virus re-infected (RI, n=15) conidia. **b.** Lung CFUs at 24h with 3×10^7^ conidia. n≥6 biologically independent biological samples from 2 independent experimental repetitions. **c.** BALF LDH levels in *A. fumigatus*-challenged mice 48 h post-inoculation with 6×10^7^ conidia. Values are presented as a percentage of LDH release from BALF cells treated with 0.1% Triton X-100 (100% cytotoxicity). n≥12 biologically independent biological samples from 3 independent experimental repetitions. **d.** Micrographs of 48 h-infected lung sections stained with hematoxylin and eosin (scale bar, 4 mm) and calcofluor white (scale bar, 50 μm). Images representative of two biologically independent experiments n=6. **e.** Morphometric analysis of lung consolidation (n = 6) at 48 hpi with 6×10^7^ conidia. **f.** Relative mRNA expression levels of proinflammatory mediators were assessed via RT-qPCR using total RNA extracted from lung homogenates at 72 hpi in mice infected with the indicated strains (Af293, n=7; VC, n=7; RI, n=6). The data are pooled from two independent experimental replicates, normalized to actin expression, and the presented values are relative to the expression levels in naïve lungs. **i.** Survival of mice challenged with 6×10^7^ Af293 or VC conidia (pretreated with ribavirin or PBS for four h) and treated with 40 mg/kg ribavirin or vehicle control (PBS) i.p. twice daily. Af293+PBS n=15, Af293+ribavirin n=15, VC+PBS n=5, VC+ ribavirin n=5. **j-k.** Pulmonary fungal burden (**j**) and mycoviral load (**k**) in mice 72 h.p.i with the indicated strains and treatments (PBS or 40 mg/kg ribavirin twice daily) were determined by qPCR. **j.** Fungal DNA was quantified per 1 µg of lung genomic DNA, using a standard curve based on fungal 18S rDNA in Af293 genomic DNA to calculate the amount of fungal DNA in the experimental samples. n = 9 for Af293+PBS, n=11for Af293+ribavirin, n=10 for VC+PBS, n=12 for VC+ribavirin. **i.** Relative expression levels of viral RdRp were assessed by qRT-PCR using the 2^−ΔCt^ method, normalized to fungal *actA* expression (reference gene). A Ct value greater than 35 or undetermined was considered indicative of viral clearance and is represented as 0. **j-k.** Data are pooled from two independent experiments. Statistical analysis: (**a, g, i**) Log-rank (Mantel-Cox). (**b-c, e-f, i**) one-way ANOVA with Tukey’s multiple comparison test. (**h**) Kruskal-Wallis test with Benjamini-Hochberg post-test, (**k**) two-tailed unpaired t test. Conidial suspensions were adjusted based on viability staining to account for differences in conidial viability prior to each experiment. Error bars indicate standard error around the mean (center).

### Antiviral Treatment Reverses the Exacerbated AfuPmV-1M-Mediated Virulence

To assess whether therapeutic intervention to block mycoviral replication could reverse the exacerbated pathology and improve the infectious outcomes, mice were challenged with Af293 or VC conidia and treated intraperitoneally with ribavirin, a nucleoside analog that targets viral replication, twice daily. Remarkably, we observed significantly improved survival in Af293-challenged mice treated with ribavirin, with a mortality rate comparable to that of virus-cured-challenged mice (Fig. 6i). Furthermore, the increased survival in the Af293-challenged, ribavirin-treated group was associated with a reduced fungal burden (Fig. 6j) and lower mycoviral load (Fig. 6k) in the lungs. Notably, ribavirin treatment showed no antifungal activity in vitro (Supplementary Fig. 8g), and survival and fungal burden in mice infected with the VC strain were not affected by ribavirin treatment (Fig. 6i-k). These data suggest that AfuPmV-1M is a viable target, and ribavirin, along with similar antiviral drugs targeting viral replication, represents a promising “antipathogenicity” treatment for virus-bearing pathogenic fungi.

## Discussion

One key characteristic of mycoviral biology is the absence of a lytic extracellular phase. As a result, mycoviruses are transmitted either vertically from one generation to the next through asexual and sexual spores, or horizontally through hyphal anastomosis ^53^. This transmission mode creates a dynamic where mycoviruses and their fungal hosts are intricately linked in terms of reproductive success. The presence of anti-mycoviral defense mechanisms within infected fungi can lead to the elimination of mycoviruses, while enhanced mycoviral virulence risks driving the fungal host to extinction. Therefore, the delicate equilibrium between mycoviral survival and the fitness of their fungal hosts plays a pivotal role in shaping the dynamics of this complex relationship. In this context, it is reasonable to anticipate the emergence of mutualistic interactions that enhance the reproductive success of both mycoviruses and their fungal hosts. Consistent with this notion, the findings of this study demonstrate a fungus-mycovirus interaction in which a dsRNA mycovirus provides a survival advantage to the fungus, particularly under environmental stress conditions and within the murine lung.

While mycoviruses have been observed to influence a wide array of phenotypes in plant fungal pathogens ^54,55^, there is limited knowledge regarding mycoviruses in human pathogenic fungi and their effects on fungal pathogenesis. In this study, we demonstrate that the mycovirus AfuPmV-1M is integral to conidiation, fitness, and virulence in *A. fumigatus*, as its absence disrupts these processes in virus-cured strains. Conidiation, a critical developmental stage tightly regulated by environmental cues, governs fitness outcomes shaped by trade-offs between growth, dispersal, and pathogenicity^56–58^. Our findings suggest that AfuPmV-1M enhances fungal fitness across both host-associated and environmental contexts without apparent fitness costs under optimal conditions. The absence of fitness trade-offs points to a highly evolved partnership between AfuPmV-1M and *A. fumigatus*, likely refined over an extended evolutionary timeframe. Such a relationship implies that potential costs of viral infection, such as metabolic burden or replication interference, have been mitigated, possibly through compensatory mutations in both fungal and viral genomes that stabilize this mutualistic interaction^59,60^. This is underscored by the consistent downregulation of the conidiation regulator BrlA in virus-cured strains, highlighting AfuPmV-1M’s adaptive integration into fungal developmental pathways (Fig. 2g). AfuPmV-1M promotes conidiation and elevates the expression of the key transcription factor *brlA*, which regulates conidiation, in both the naturally virus-infected and congenic re-infected strains, compared to the congenic virus-cured strain (Fig. 2 and Supplementary Fig. 3). These findings are consistent with previous studies on closely related polymycoviruses, where naturally virus-infected strains exhibited darker pigmented colonies, increased conidia production, and elevated BrlA expression compared to their congenic virus-cured counterparts^11,61^. Importantly, BrlA positively regulates 13 BGCs involved in the synthesis of secondary metabolites specific to vegetative growth and asexual development, including DHN-melanin, a well-established virulence factor in *A. fumigatus*^25^. Consistent with this and in line with metabolic control principles—where rate-limiting steps in biosynthetic pathways often occur at the initial substrate-to-product conversion or final product formation step—we observed transcriptional upregulation of the first (*pksP*) and last (*arb2*) genes in the DHN melanin biosynthesis pathway in the virus-infected strains (Fig. 2h). This suggests an increase in DHN melanin production in these strains. The contribution of DHN-melanin to *A. fumigatus* virulence is partly attributed to its cytoprotective effect, which aids survival in hostile environments, particularly under oxidative stress conditions^62,63^. Indeed, our data strongly suggest that AfuPmV-1M confers survival advantages to its fungal host under various stress conditions and within mammalian lungs (Fig. 3). We found that eliminating AfuPmV-1M from a naturally infected strain reduced the expression of key regulators of DHN melanin production, correlating with diminished fungal fitness under oxidative and heat stress. This reduction also corresponded with decreased survival in the lung environment and attenuated virulence, suggesting that impaired melanin regulation may partially explain the observed reductions in fitness and pathogenicity. Intriguingly, the enhanced fitness conferred by AfuPmV-1M was most striking in swollen conidia under stress, while virus-infected resting conidia displayed moderately improved resilience against oxidative, heat, pH, and osmotic challenges compared to their virus-cured counterparts (Fig. 3a, Supplementary Fig. 4). The upregulation of AfuPmV-1M RdRP in swollen conidia, relative to resting conidia, further ties AfuPmV-1M to the enhanced fitness phenotype observed under stress in swollen virus-infected conidia compared to virus-cured counterparts (Fig. 3b). This finding may explain discrepancies with prior studies reporting reduced fitness in virus-infected strains under oxidative stress, a trend likely tied to their focus on resting conidia^12,13^. In contrast, our data indicate that virus-infected resting conidia exhibit moderately improved resilience to oxidative, heat, pH, and osmotic stresses compared to virus-cured counterparts, though this effect is less marked than in swollen conidia. Notably, both our study and these earlier studies utilized the naturally AfuPmV-1M-infected *A. fumigatus* Af293 strain and its congenic cured derivative, suggesting that variations in experimental conditions or stress protocols may account for the contrasting phenotypes observed. Specifically, prior work tested oxidative stress resistance at 32.6 mM H₂O₂ or higher concentrations, compared to 5 mM H₂O₂ in our study—a concentration more aligned with physiological levels encountered in the lung environment during phagocyte interactions^64,65^. This higher concentration could amplify stress responses, potentially masking the moderate resilience we observed in virus-infected resting conidia under our milder conditions. Unlike our data (Supplementary Fig. 2) and that of Kanhayuwa et al.^11^, which show comparable growth and germination between virus-infected and virus-cured strains under optimal conditions, these studies reported attenuated growth and germination in the virus-infected strain under similar conditions ^12,13^. This variation may reflect differences in basal strain physiology or subtle environmental factors not captured in our assays, suggesting that AfuPmV-1M’s effects are highly context-dependent, manifesting most prominently in swollen conidia under host-relevant stress levels and to a lesser extent in resting conidia, rather than under optimal growth conditions. Further supporting the integration of AfuPmV-1M with fungal fitness, we observed differential expression of all five mycoviral ORFs under various stress conditions in the naturally infected Af293 strain (Fig. 3c). Notably, all ORFs were upregulated under stress and during infection, correlating with a cytoprotective phenotype that enhances fungal resilience. This contrasts with findings by Takahashi-Nakaguchi et al. [14], who reported increased susceptibility to stresses upon heterologous expression of individual AfuPmV-1M ORFs in a naïve *A. fumigatus* KU strain^14^. We propose that this discrepancy arises from differences in the immune state of the fungal host. In our system, Af293 has co-evolved with AfuPmV-1M, likely establishing a symbiotic relationship where cellular processes—including fungal antiviral immune responses—are co-regulated to bolster both fungal and mycoviral survival. Conversely, Takahashi-Nakaguchi et al. overexpressed isolated ORFs outside the context of an intact virus in the KU strain, a derivative of *A. fumigatus* CEA17 lacking the ku80 gene^66^ and naïve to AfuPmV-1M. This strain likely lacks the co-evolved immune state of Af293, which may explain its heightened susceptibility when expressing mycoviral ORFs. We hypothesize that the immune status of the fungal host is a critical determinant of AfuPmV-1M’s impact on fitness: in Af293, mycoviral replication is synchronized with host physiology, conferring a fitness advantage, whereas in a naïve strain, viral replication may not be supported or could even be detrimental to the fungal host. This host-virus interplay underscores the context-dependent nature of mycoviral effects and highlights the need to consider fungal immune adaptation in interpreting their biological roles.

To elucidate the molecular mechanisms driving AfuPmV-1M’s context-specific fitness effects, we performed a proteome analysis to identify key proteins and pathways involved. Our proteome analysis revealed that AfuPmV-1M infection upregulates proteins involved in RNA metabolism, notably Brf1 and RNA Polymerase III (Pol III), which we further supported by RT-qPCR showing increased expression of Pol III targets, including tRNAs and regulatory non-coding RNAs (Fig. 4a-d). This heightened RNA metabolism suggests a dual viral strategy: AfuPmV-1M may hijack the fungal RNA machinery to enhance transcription of viral RNA or host RNAs critical for replication, leveraging increased tRNA pools for protein synthesis, as seen in other RNA viruses ^67,68^; concurrently, it could bolster fungal resilience under stress, supporting host survival and viral persistence, consistent with mycovirus-induced fitness enhancements. Strikingly, we also observed upregulation of proteins known to localize to stress granules (SGs)—cytoplasmic assemblies that regulate RNA stability and translation during stress (Fig. 4a, e). Using a fungal stress granule reporter, we confirmed that the virus-infected strain forms significantly more stress granules than its virus-cured counterpart, indicating a robust stress response (Fig. 4f-h). These findings suggest that AfuPmV-1M links RNA metabolism to stress granule (SG) formation, akin to previous studies demonstrating RNA excess as a driver of SG assembly ^69,70^. Thus, upregulated RNA production may both fuel viral replication and promote granule formation to enhance cytoprotection under environmental and host-relevant conditions. In light of our proteome data, we propose that AfuPmV-1M exploits fungal survival pathways by inducing transcriptional and translational modifications that modulate components such as stress granules, enhancing fungal resilience and viral replication under impending harsh conditions. This modulation, in turn, influences the virulence of the fungal host. Specifically, we suggest that chronic AfuPmV-1M infection imposes a mild, non-lethal cellular stress that primes *A. fumigatus* for subsequent lethal challenges, conferring’cross-tolerance’ against diverse insults, including oxidative and heat stress— common stressors in both environmental niches and host interactions. This priming effect mirrors a hormetic adaptation observed in cancer cells, where stress granules induced by mild stress precondition cells for severe stresses, enhancing survival and stress resistance ^71^; although these SG-mediated priming studies focus on mammalian cells, microbial priming was proved a common defense strategy ^72^. Our model is further supported by the observation that Thioredoxin 1 (Trx1), a key enzyme protecting cells from oxidative damage ^49^, and a stress granule component essential for their protective function ^45^, is upregulated in AfuPmV-1M-infected strains compared to their virus-cured counterparts (Fig. 4). Collectively, these findings indicate that infection by AfuPmV-1M orchestrates a sophisticated stress response strategy, balancing fungal survival while potentially sustaining its own propagation.

Based on current insights into fungal pathogenicity, *A. fumigatus* strains with enhanced resistance to oxidative stress are predicted to exhibit improved survival within the pulmonary environment, potentially resulting in more severe disease^51,73–75^. Consistent with this hypothesis, our data demonstrate that the virus-cured strain, which displays greater susceptibility to various stresses compared to its mycovirus-infected counterparts, exhibits reduced virulence across three immunocompetent murine strains (C57BL/6JOlaHsd, C57BL/6J, ICR) and in a neutropenic IPA model, despite maintaining comparable growth rates under standard laboratory conditions (Fig. 6a–c, Supplementary Fig8. a, c–f). As noted earlier, this cytoprotective effect is particularly evident in the swollen conidial state, a morphologically critical stage for studying *A. fumigatus* as an inhaled environmental mold pathogen. Following inhalation, conidia undergo a swelling process in the airways that exposes fungal pathogen-associated molecular patterns (PAMPs), triggering recruitment of immune cells; consequently, their initial encounter with host phagocytes, specifically neutrophils, occurs in this swollen state^76,77^. This swollen conidial state thus represents a key interface for host-pathogen interactions in our system, highlighting its relevance to mycovirus-mediated virulence modulation. In contrast, Takahashi-Nakaguchi et al. (2020) reported reduced virulence of a mycovirus-infected *A. fumigatus* strain compared to its virus-cured counterpart in a neutropenic mouse IPA model, a finding that sharply contrasts with our observation of enhanced virulence in both immunocompetent and neutropenic mice^14^.

This discrepancy likely arises from experimental differences. For instance, our study utilized C57BL/6J mice, a widely used inbred strain in fungal infection models, whereas Takahashi-Nakaguchi et al. employed ICR mice, an outbred strain. Additionally, the fungal inoculum differed substantially between the studies: we administered 5 × 10^3^ conidia per mouse, compared to 5 × 10^7^ conidia per mouse in their neutropenic murine model. Their use of antibiotic treatment in the ICR mice may have influenced immune responses, potentially contributing to the observed differences in virulence outcomes. Moreover, while Takahashi-Nakaguchi et al. observed reduced survival of the virus-infected strain in a macrophage cell line using resting conidia, we demonstrated enhanced survival of virus-infected conidia in vivo when naturally swollen and phagocytosed by neutrophils at 24 h post-infection, and ex vivo following phagocytosis of artificially pre-swollen conidia by human-derived neutrophils (Fig. 5). This contrast further emphasizes the physiological relevance of the swollen conidial state in our system, a finding consistently observed across two virus-infected strains (the naturally infected Af293 and a congenic re-infected strain), tested in three immunocompetent and one neutropenic murine model of IPA.

Although immunocompromised individuals constitute the primary population at risk for invasive aspergillosis, the use of an immunocompetent mouse model in our study offers critical insights into the full spectrum of host immune responses and fungal survival strategies under conditions mimicking early infection stages in healthy hosts, potentially informing preventive strategies and initial pathogenesis mechanisms^78,79^. In our system, where *A. fumigatus* harbors a mycovirus, the mammalian immune system may target both the fungal pathogen and the mycovirus itself; thus, an immunocompetent murine model enables examination of a healthy immune response to the fungus in the context of a mycoviral infection during the early infection stages following the initial encounter with phagocytes.

This approach complements neutropenic models by uncovering mycovirus-mediated effects on virulence that might otherwise be obscured in the absence of intact immunity.

Importantly, we introduce a pioneering therapeutic strategy by targeting an endogenous mycovirus to attenuate the pathogenicity of its fungal host, a novel approach demonstrated here for the first time in vivo. Specifically, we utilized the antiviral nucleoside analog ribavirin to target AfuPmV-1M during IPA in a murine model, rescuing *Aspergillus*-infected mice by reducing the virulence of the mycovirus-infected strain to levels comparable to its virus-cured counterpart, an effect correlated with diminished fungal burden and viral load (Fig. 6i–k). We present this as a proof-of-concept for mycoviruses as modulators of fungal virulence, with ribavirin reducing fitness by targeting AfuPmV-1M replication, rather than a direct endorsement of antivirals as adjunct therapy for invasive aspergillosis (IA), given that human IA predominantly occurs in immunocompromised hosts—a context distinct from our high-dose immunocompetent model. Unlike prior studies that employed antiviral drugs solely to generate mycovirus-cured strains ex vivo, our in vivo intervention highlights ribavirin’s potential as an ‘antipathogenicity’ treatment, directly mitigating disease severity in an active infection. To our knowledge, this represents the first demonstration of such a therapeutic approach not only for mycovirus-bearing fungi but also in the broader context of eukaryotic pathogens and their endogenous viruses, such as *Leishmania*, where the well-studied Leishmania RNA virus-1 (LRV-1) enhances virulence, yet antiviral drugs have been limited to generating virus-cured congenic lines ex vivo ^80^. This repurposing of existing antiviral drugs offers a transformative avenue for managing invasive fungal infections associated with mycoviruses, with broader implications for targeting other infectious diseases where pathogenicity may be exacerbated by similar endogenous viruses. Our cyclophosphamide-induced neutropenic model confirms that the AfuPmV-1M-mediated virulence phenotype persists in an immunocompromised setting, suggesting that ribavirin and similar replication-targeting antivirals could be effective in such contexts, though this requires future investigation beyond our current scope. By leveraging clinically approved antivirals, this strategy could accelerate the development of personalized, effective therapeutic regimens that account for the presence of fungal viruses, ultimately improving patient outcomes and reducing healthcare costs associated with recalcitrant fungal infections.

Our findings reinforce the multifactorial nature of fungal virulence, revealing it as a dynamic interplay of fungal traits, host responses, and viral ‘hitchhikers’ that may act as critical backseat drivers in disease progression. Although this study focuses on a reference strain of *A. fumigatus* from clinical origin, expanding to additional environmental and clinical mycovirus-bearing isolates in future research could establish this as a widespread phenomenon, where environmental pressures—both biotic and abiotic—select for mycovirus-fungal partnerships that enhance fitness and stress resistance, conferring competitive advantages in niches like compost or the mammalian lung. Such mycovirus-derived traits may drive fungal host adaptation and the emergence of novel pathogenic strains from environmental reservoirs. Most significantly, by coupling mycoviruses with virulence and demonstrating their targetability with existing therapeutics, our work lays the foundation for expanding diagnostic and therapeutic frameworks beyond traditional fungal species and strain identification. Incorporating the detection and targeting of endogenous viruses that profoundly influence virulence could revolutionize the management of fungal pathogens, offering a paradigm shift in both clinical diagnostics and treatment strategies.

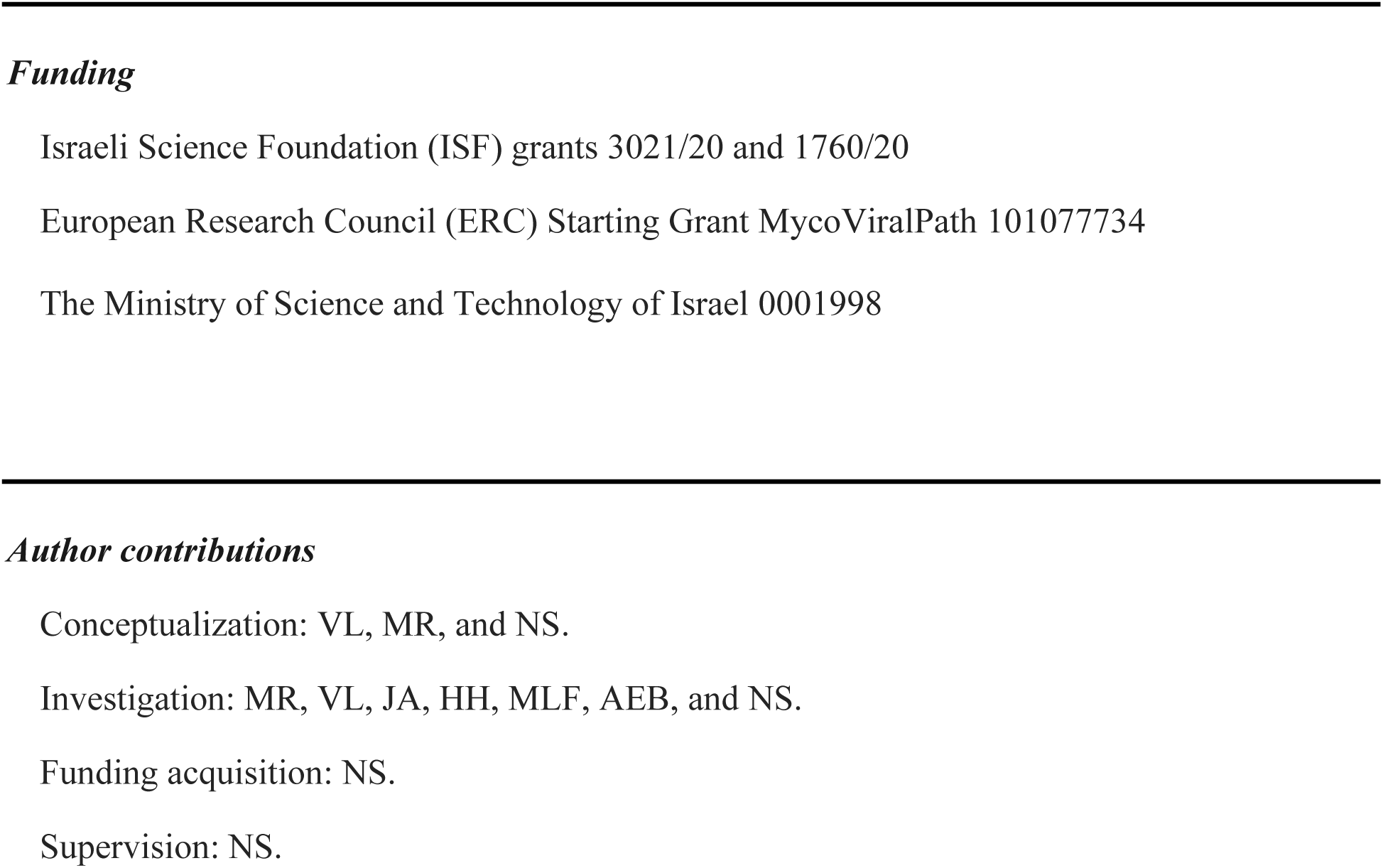
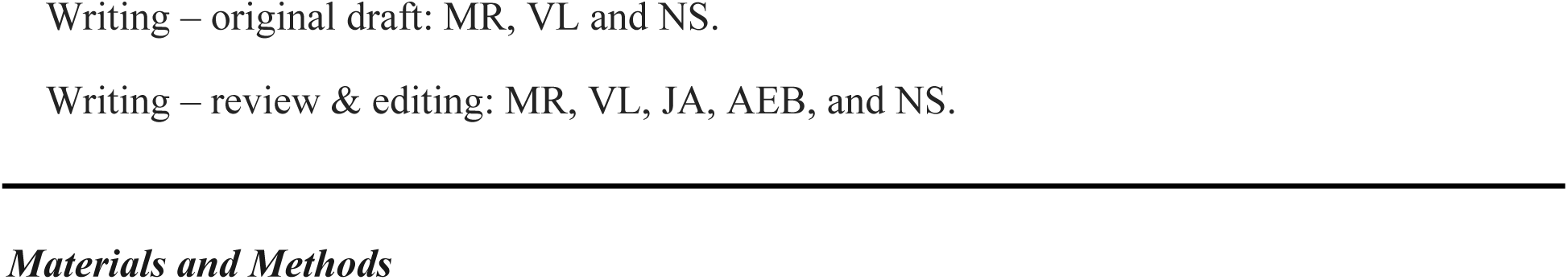

### Fungal strains and growth conditions

The naturally Afupmv-1M-infected *Aspergillus fumigatus* strain *Af293* was obtained from Dr. Tobias Hohl and kept in our laboratory for over four years. Virus-cured strains (VC) were generated from *Af293* through treatment with either two cycles of 1mM and 10 mM of the protein synthesis inhibitor cycloheximide or 0.1 mg/ml of the nucleoside analog ribavirin, as previously described ^22,81^. A congenic re-infected strain (RI) was obtained by PEG-mediated transfection of 50 µg of purified AfuPmV-1M dsRNA, extracted from the parental *Af293* strain into the cycloheximide-cured strain (VC) as previously described ^81^. All strains were preserved, cultured, and collected for experimentation as previously described ^27,51^. For the purification of viral particles and viral dsRNA, fungal cultures were grown in 250 ml of GMM broth with continuous agitation at 130 rpm, at 37°C for a period of 5-7 days.

Subsequently, the mycelium was harvested with Miracloth, snap-frozen with liquid nitrogen, and stored at-80°C until subsequent processing.

### Purification of viral particles

The viral particle purification process was conducted as previously described ^11^ with minor adaptations. Approximately 50 grams 6-day-old mycelia from *A. fumigatus Af293* were homogenized in a 3-fold volume (wt/vol) of 0.1 M sodium phosphate solution at pH 7.4, supplemented with 0.2 M KCl and 0.5% mercaptoethanol using a manual homogenizer (Hangzhou Miu Instruments Co. LTD, type MT-30K). The resulting homogenate was clarified with an equal volume of chloroform followed by centrifugation at 10,000×*g* for 20 min at 4°C. The virus-containing supernatant was mixed with 10% (wt/vol) PEG-6000 and 0.6 M NaCl, and incubated overnight at 4°C with constant stirring. The following day, the suspension underwent low-speed centrifugation at 10,000×*g* for 20 min. The resulting precipitate, including viral particles, was resuspended in 60 mL of TE buffer and clarified through additional low-speed centrifugation. Any excessive PEG contamination pellet was discarded, while the remaining supernatant, which contained the virus, was subjected to ultracentrifugation at 105,000×*g* for 90 min at 4°C. The pellet enriched with virus-like particles (VLPs) was resuspended in 1 mL of TE buffer.

### RNA extraction, RT-PCR, and qRT-PCR

Mycelia from liquid shaking cultures were flash-frozen, powdered in liquid nitrogen, and homogenized in Trizol (300 mg mycelia in 1 ml Trizol). For RNA extraction from conidia, conidia were collected from 3 days-old plates and resuspended and homogenized in 1 ml of Trizol. Mycelial and conidial RNA was extracted as previously described ^11^. 10 µg of RNA was DNase-treated with TURBO DNA-free (Invitrogen™ # AM1907), according to the manufacturer’s instruction. cDNA was obtained using High-Capacity cDNA Reverse Transcription Kit (Applied Biosystems™ # 4368814). For the expression analysis of viral genes, pretreated with 15% DMSO (v/v) for 10 min at 95°C has been used to denature dsRNA prior to the reverse transcription ^82^. The presence of targeted genes was confirmed by PCR amplification using Promega GoTaq Green Master Mix. The primers for the individual genes are listed in Supplementary Material (Table S1). qRT-PCR was performed using the Fast SYBR® Green Master Mix (Applied Biosystems™, # 4385612) and data was collected in a StepOne Plus Real Time PCR System (Thermo Scientific). The concentration of each primer pair was optimized prior to the efficiency curve reaction. Only primers with amplification efficiency ranging from 95–105% were used, according to reference^83^.

Nontemplate controls (NTC) were used to confirm the elimination of contaminating DNA in every run. A melt curve analysis was performed after the PCR was complete to confirm the absence of nonspecific amplification products. The fold change in mRNA abundance was calculated using 2^−ΔΔCt^ ^84^ and all the values were normalized to the expression of the *A. fumigatus* 18S or *actA* for measurement of fungal and mycoviral genes or to *actβ* for measurements of murine genes. Fungal burden was quantified by qPCR of the 18S rDNA region, as previously described ^85^.

### Isolation of dsRNA by LiCl precipitation

Total RNA was extracted from *A. fumigatus* 250 ml liquid culture grown at 37°C until saturation, approximately 5 to 6 days. ssRNA/dsRNA fractionation was performed as previously described ^86^. To visualize dsRNA, 1 µg was pretreated with RNase-free DNase1 (RBC Bioscience, cat. #9680DN050) and S1 nuclease (Thermo Scientific™, cat. #EN0321) according to the manufacturer’s instructions and electrophoresed on 1% agarose gel stained with ethidium bromide.

### Whole-genome DNA sequencing and analysis

*A. fumigatus* genomic DNA was extracted as previously described^87^. Illumina sequencing libraries were prepared by Seqcenter (SeqCenter LLC, PA, USA) using the tagmentation based and PCR-based Illumina DNA Prep kit and custom IDT 10bp unique dual indices (UDI) with a target insert size of 320 bp. Illumina sequencing was performed on an Illumina NovaSeq 6000 sequencer, producing 2×151bp paired-end reads. Demultiplexing, quality control and adapter trimming was performed with bclconvert1 (v4.1.5). The raw FASTQ reads were preprocessed using fastp v0.23.4^88^, removing Illumina adaptor sequences and sequences with low quality scores (Phred score < 30). The quality of the sequences before and after quality control was assessed using FastQC v0.11.9 (Babraham Institute). Reference-guided genome analysis was conducted on high-quality sequence reads by aligning them to the *A. fumigatus* Af293 reference genome (FungiDB release 66) using Bwa-mem2 v2.2.1^89^, followed by conversion to BAM files using samtools v1.16.1^90^. PCR duplicates were identified and marked using the MarkDuplicatesSpark implementation within GATK v4.4.0.0^91^. All WGS samples analyzed exhibited a genome coverage of >10-fold after mapping, with >95% of reads aligning to the reference genome. Subsequently, deduplicated BAM files were recalibrated using GATK BaseRecalibrator ApplyBSQ, using an internal dataset of known variable sites derived those cataloged in FungiDB, release 39. Variant calling was carried out using GATK Haplotypecaller v4.4.0.0 with the ploidy set to 1 followed by joint calling of all samples. The resulting variants were hard-filtered using the specific parameters:’QD < 3.0’,’MQ < 40.0’,’FS > 60.0’,’MQRankSum <-10.0’,’ReadPosRankSum <-5.0’ for SNPs, and’QD < 3.0’,’FS > 200.0’,’ReadPosRankSum <-20.0’ for INDELs. Additionally, variants were identified using the Platypus variant caller 0.8.1.2^92^, with filter parameters set to a quality filter of 30, a depth filter threshold of 10, and an allele frequency filter threshold of 0.01. Following variant calling, the VCF files from HaplotypeCaller and Platypus were merged using bcftools^90^ isec v1.17. Next, background variants that were also detected in our lab copy of AF293 were subtracted from the VC and RI samples using bcftools. Finally, the variants were annotated using SnpEff v4.3^92^ using a cutoff of 1,000 bp for upstream and downstream regions. Identified SNPs underwent visual inspection of alignments using IGV version 2.17.0^93^ to detect false positive variant calls that may have evaded automated filters. Samples were screened for structural variants, including copy number variations, and aneuploidies using controlFREEC v11.6^94^ and DELLY v1.1.8^95^ and only structural variants found by both tools were taken as true SVs. For controlFREEC, sequencing data from the lab copy of AF293 was input as the control and the additional parameters of ploidy=1, minExpectedGC=0.35, and maxExpectedGC=0.60 were used.

Significance testing was performed using the tool provided script and CNVs were filtered using Bonferroni-corrected Wilcoxon Rank Sum Test p < 0.05. DELLY was run with the ploidy set to 1 and using a mappability file created using Dicey v0.2.6. Low quality predictions were removed by filtering out SVs flagged by DELLY as ‘LOW QUALITY’. Finally, any CNVs identified by both controlFREEC and DELLY were manually inspected in IGV v2.18.2.

### Stress assays

10^6^ conidia/ml conidia of *A. fumigatus Af293*, VC, and RI strains were incubated in GMM broth for 4 h at 37°C, to obtain swollen conidia. Subsequently, conidia were subjected to stressors including 5mM H_2_O_2_, pH 4.5, pH 8.5, 50°C, and 1 M Sorbitol for 4 h at 37°C, 130rpm. Conidial viability was assessed by colony forming units (CFU). To evaluate growth inhibition of *A. fumigatus* strains 10^3^ conidia were inoculated onto GMM agar plates under control conditions or at pH 4.5, pH 8.5 and 1M Sorbitol, or 50°C. Diameter of the colony was measured on the 5^th^ day of incubation.

### Biofilm formation assay

Briefly, 10^6^, 10^5^, or 10^4^ spores in 1% GMM broth were seeded per well in a 24-well microtiter plate with or without voriconazole at the indicated concentration were incubated at 37°C for 48 h. To determine the biofilm formation ability of *A. fumigatus* strains, we conducted the crystal violet (CV) assay as previously described ^96^ with minor modifications. 1 mL of GMM broth with and without the voriconazole was inoculated with 10^6^, 10^5^, and 10^4^ conidia in a 24-well microtiter plate, and incubated at 37°C for 48 h. The plate was then washed twice with PBS to eliminate free-floating cells, and the biofilms were stained with 1 mL of 0.1% crystal violet solution for 10 min at room temperature. Biomass was washed with PBS at least 5 times and air-dried, followed by incubation with 1 mL of 100% acetic acid, slightly shaken at room temperature for 30 min to elute the crystal violet, and the absorbance was measured at 470 nm using a Tecan Infinite Pro Plate reader.

### Phagocytosis assay

Human blood was obtained from healthy volunteers, with informed consent, in compliance with the guidelines of the Hebrew University of Jerusalem Research Ethics Committee. Human neutrophils were harvested and *Acanthamoeba castellanii* were maintained as previously described ^97,98^. For the phagocytosis and survival assay with either amoeba or human neutrophils, 1×10^6^ phagocytes were plated in a 24-well plate containing 1 ml of supplemented DMEM (human neutrophils) or PYG (amoeba) medium and pre-incubated for 1 h letting amoebal cells attach to the surface. The cell walls of swollen conidia (4 h in GMM at 37°C) were fluorescently labeled with Alexafluor 488-conjugated streptavidin as previously described ^50,51^. 1×10^6^ labeled *A. fumigatus* conidia were inoculated into the neutrophil and amoeba culture and incubated for 4 h at 37°C under 5% CO2 (neutrophils) or 30°C (*A. castellanii)*. Phagocytosis was evaluated by flow cytometry. The phagocytes’ population was gated by size, and the AF488 ^+^ population represents conidial uptake. Conidial viability was evaluated by colony forming units (CFUs).

### Conidiation and germination assays

Conidiation was quantified by harvesting conidia from 4 days old GMM agar plates that were inoculated with 2×10^5^ conidia. The conidial concentration was determined by counting with a hemocytometer. To determine the germination rate, conidia were collected from 5 days-old GMM plates, adjusted to 5×10^5^ conidia per ml in GMM broth and inoculated in 8 wells microscopy chambered slide at 37°C. Images were captured using an inverted motorized Nikon fluorescence microscope (Eclipse Ti2) in time-lapse mode, with 30 min frame over a period of 24 h.

### Quantification of conidia during asexual development

Synchronized asexual differentiation was conducted as described previously ^99^. Briefly, 1×10^8^ conidia of the virus-infected (Af293 and RI), or virus-cured strains were inoculated in 100 ml of liquid minimal medium incubated at 37°C (200 rpm) for 16 h. Mycelia from the submerged cultures (5 g [wet weight]) were then harvested by filtration and flash frozen in liquid nitrogen (t0 control samples) or transferred to a solid, complete medium and further incubated at 37°C (for 6, 12, and 24 h). Samples were collected at each time point flash frozen in liquid nitrogen and stored at –80°C until they were used for RNA extractions.

Conidial counts for each strain subjected to synchronized asexual differentiation were conducted by sampling 0.5 g of mycelium at 6, 12, and 24 h after the induction of conidiation. The conidia were collected in a 0.01% Tween 20 solution and counted using a Neubauer chamber. The experiments were repeated three times.

### Inspection of fungal lawn and visualization of conidiophores

For evaluation of fungal lawn morphology, 200 ul of 10^6^ conidia/ml was spread onto GMM agar plates and incubated for 7 days at 37°C. 10 ul of 2×10^6^ conidia were inoculated into agar slices and covered by a sterile coverslip. After 3 days after inoculation, glasses with attached conidiophores were gently detached from the agar slice, placed on microscopy glass with lactophenol blue, and sealed by polish. Conidiophores were visualized by ×100 microscopy (Nikon, Eclipse Ti2), photos were captured and conidia on individual conidiophores were counted.

### Mice, animal care, and ethics statement

6-8 weeks-old C57BL/6JOlaHsd and ICR mice were purchased from Envigo, Israel. C57BL/6J mice were originally obtained from The Jackson Laboratory (strain 000664) and bred under specific pathogen-free conditions at our mouse facility at the Hebrew University of Jerusalem. All animal experiments were conducted with sex-and age-matched mice and are compatible with the standards for care and use of laboratory animals. The research has been approved by the Hebrew University of Jerusalem Institutional Animal Care and Use Committee (IACUC) (protocol number MD-20-16072-5). The Hebrew University of Jerusalem is accredited by the NIH and by AAALAC to perform experiments on laboratory animals (NIH approval number: OPRR-A01-5011).

### Murine models of IPA

The virulence of the *A. fumigatus* strains was assessed in two murine models of IPA, immunocompetent and the chemotherapeutic models. For the chemotherapeutic murine model, C57BL/6J mice 8 to 10 weeks old, were immunosuppressed with intraperitoneal (i.p.) injections of cyclophosphamide (Western Medical Supply, Inc.) at 250 mg/kg of body weight 48 h before fungal inoculation and 72 h after fungal inoculation. Mice were infected via the intranasal route as described in ^27^. In assays where labeled conidia were used, the conidia were labeled as previously described ^50^. BAL and lung suspensions were prepared for flow cytometry as described in ^51^. Briefly, lung digest and, if applicable, BAL cells were enumerated and stained with the following Abs: anti-Ly6C (clone AL-21), anti-Ly6G (clone 1A8), anti-CD11b (clone M1/70), anti-CD11c (clone HL3), anti-CD45.2 (clone 104), anti-MHC class II (clone M5/114.15.2), anti-Siglec F (clone E50-2440) and anti-CD103 (clone 2E7). Neutrophils were identified as CD45^+^CD11b^+^Ly6C^lo^Ly6G^+^ cells, inflammatory monocytes as CD45^+^CD11b^+^Ly6C^hi^Ly6G^−^ cells, Mo-DCs as CD45^+^CD11c^+^MHC class II^+^ CD11b^+^CD103^-^SiglecF^-^ cells, and alveolar macrophages as SSC^hi^CD45^+^CD11b^+^CD11c^+^ SiglecF^+^ bronchoalveolar lavage (BAL) fluid cells. Flow cytometry data was collected on a NovoCyte Quanteon (Agilent Technologies Inc) flow cytometer and analyzed on NovoExpress version 1.6.1 (Agilent Technologies Inc). Neutrophils containing ingested labeled conidia were sorted using the BD FACSAria™ III (BD Biosciences, San Jose, CA). BAL Lactate dehydrogenase (LDH) levels were measured with a CytoTox 96 Non-Radioactive Cytotoxicity Assay (Promega, Madison, WI).

Perfused murine lungs were homogenized in 2 mL of PBS, 0.025% Tween-20 for colony-forming units (CFUs) and ELISA. For histology, perfused lungs were fixed in 10% neutral-buffered formalin, embedded in paraffin, sectioned in 4 μm slices, and stained with hematoxylin & eosin (H&E) or calcofluor white (MP Biomedicals). Slides were reviewed and evaluated in a blinded manner. Whole section images were digitally scanned using a Nanozoomer S360 slides scanner.

### Liquid chromatography–MS/MS

Liquid chromatography–MS/MS was performed at The De Botton Protein Profiling institute of the Nancy and Stephen Grand Israel National Center for Personalized Medicine, Weizmann Institute of Science. 10^6^ conidia per ml were cultured in 75 ml of GMM broth for 18 h, followed by a 4-hour oxidative stress challenge with 8 mM of H_2_O_2_. The resulting mycelium was ground in liquid nitrogen and then resuspended in a 5% SDS buffer. The samples underwent in-solution tryptic digestion using the S-trap protocol developed by Protifi. The resulting peptides were subjected to analysis using nanoflow liquid chromatography (nanoElute) coupled to a high-resolution, high-mass-accuracy mass spectrometry system (Bruker Tims TOF Pro). During analysis, the samples were randomly ordered in discovery mode. The raw data was processed with FragPipe v17.1. The data was searched with the MSFragger search engine against the NCBI *Aspergillus fumigatus* Af293 proteome, appended with common lab protein contaminants. Carbamidomethyl was specified as a fixed modification. Oxidation of methionine residues and protein N-terminal acetylation were specified as a variable modification. The quantitative comparisons were calculated using Perseus v1.6.0.7. Decoy hits were filtered out and only proteins detected in at least 2 replicates of at least one experimental group were kept.

## Statistics and reproducibility

All statistical analysis was performed in GraphPad Prism version 8.0.2. Unless otherwise noted, all statistical analyses were performed with at least three biologically independent samples. All images are representative of a minimum of three biologically independent samples that represent a minimum of three independent experimentations unless otherwise noted. For comparisons between two groups two-tailed unpaired t-tests were performed. For comparisons between greater than two groups One-Way ANOVA with Tukey, Sidak, or Dunnett post tests for multiple comparisons were performed. All error bars indicate standard error and are centered around the mean.

## Data Availability Statement

The genomic sequencing data for the cycloheximide-cured and the re-infected strains, the ribavirin-cured strain, and the parental AF293strain been deposited under BioProject identifier <u>PRJNA1082383</u>. All strains and any other data used to support these findings will be made available upon reasonable request to the corresponding author.

## Supplementary Figures

**Figure S1.**
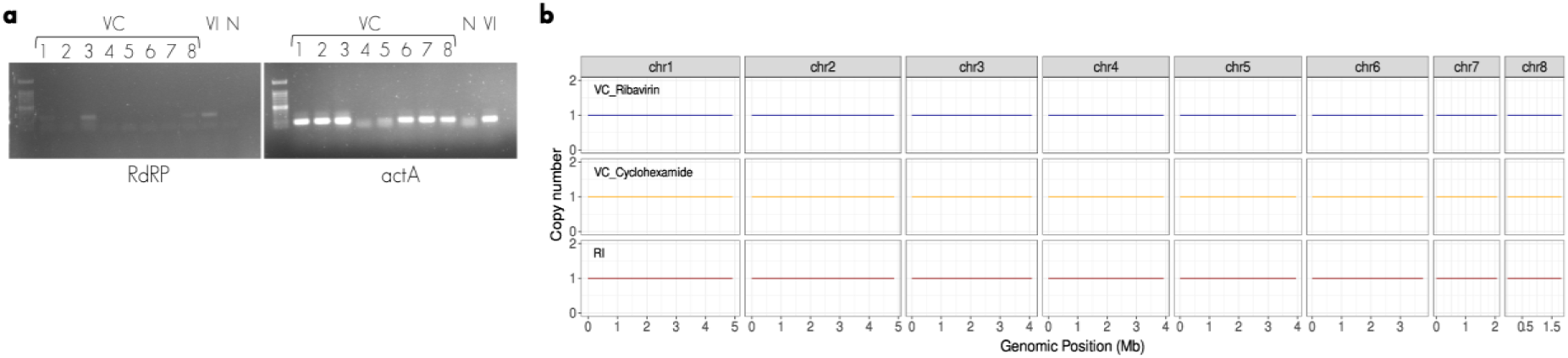
Generation of congenic ribavirin virus-cured strains and whole genome sequencing. **a.** Reverse-transcription PCR (RT-PCR) amplification of a 276 bp segment from Afupmv-1M RdRP in total RNA extracted from Af293 (VI) and ribavirin virus-cured (VC). N-negative control. **b**. Genomic comparisons of *A. fumigatus* virus-infected, virus-cured and re-infected strains compared to the reference strain Af293.

**Figure S2.**
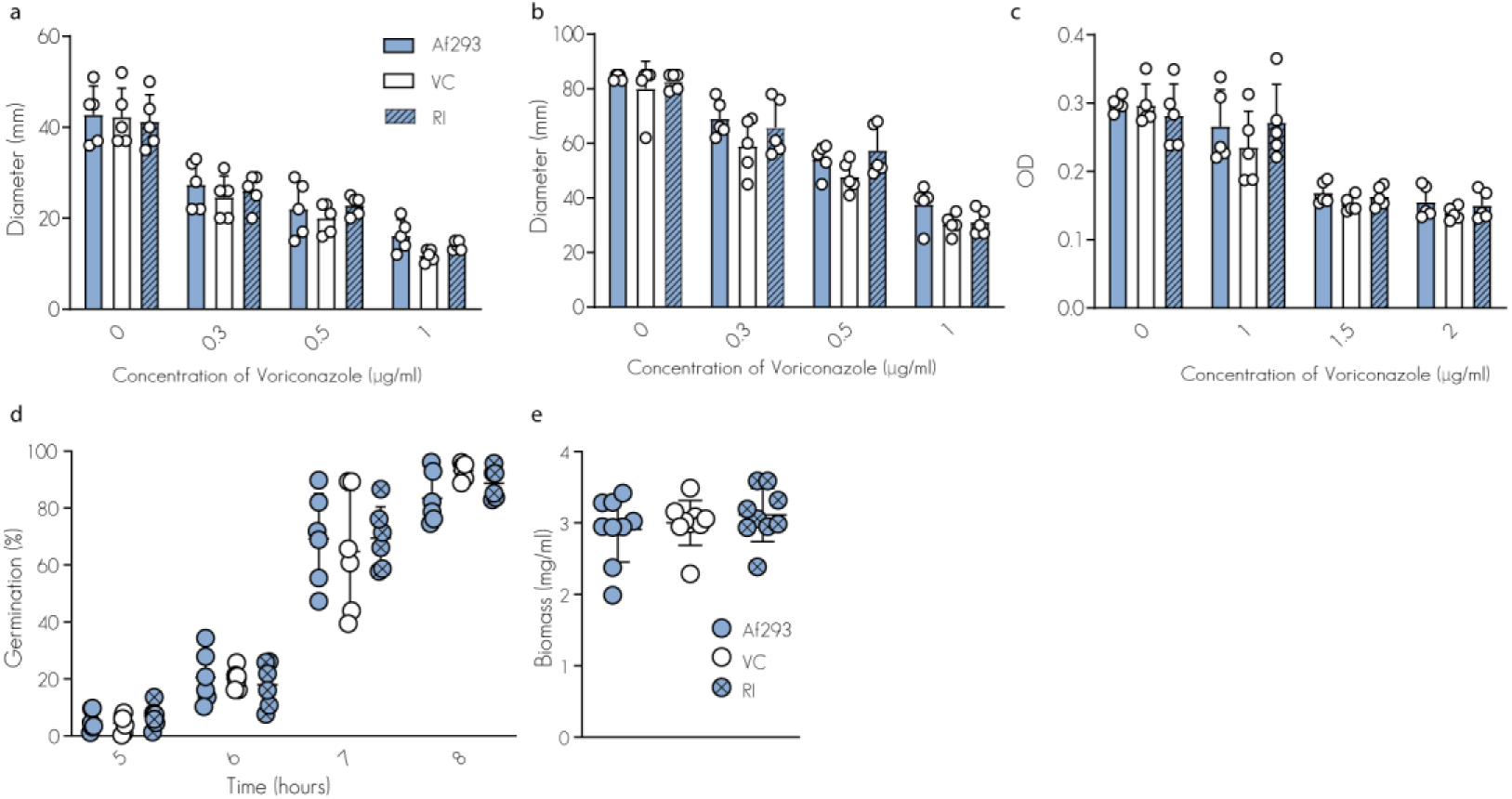
Characterization of the Afupmv-1M-cured and re-infected strains. Radial growth of Af293, cycloheximide-cured, and the re-infected strains on GMM plates amended with Voriconazole three days (**a**) and seven days (**b**) post-inoculation. **c**. Biofilm formation on a plastic surface in the presence of Voriconazole after 48 h of incubation at 37°C. **d**. Germination rate of conidia. **e**. Fungal biomass production over 24 h of incubation. Conidial suspensions were adjusted based on viability staining to account for differences in conidial viability before each experiment. The data are derived from a minimum of three biologically independent experiments. Error bars indicate standard error around the mean (center). Statistical analysis: one-way ANOVA with Tukey’s multiple comparison test.

**Figure S3.**
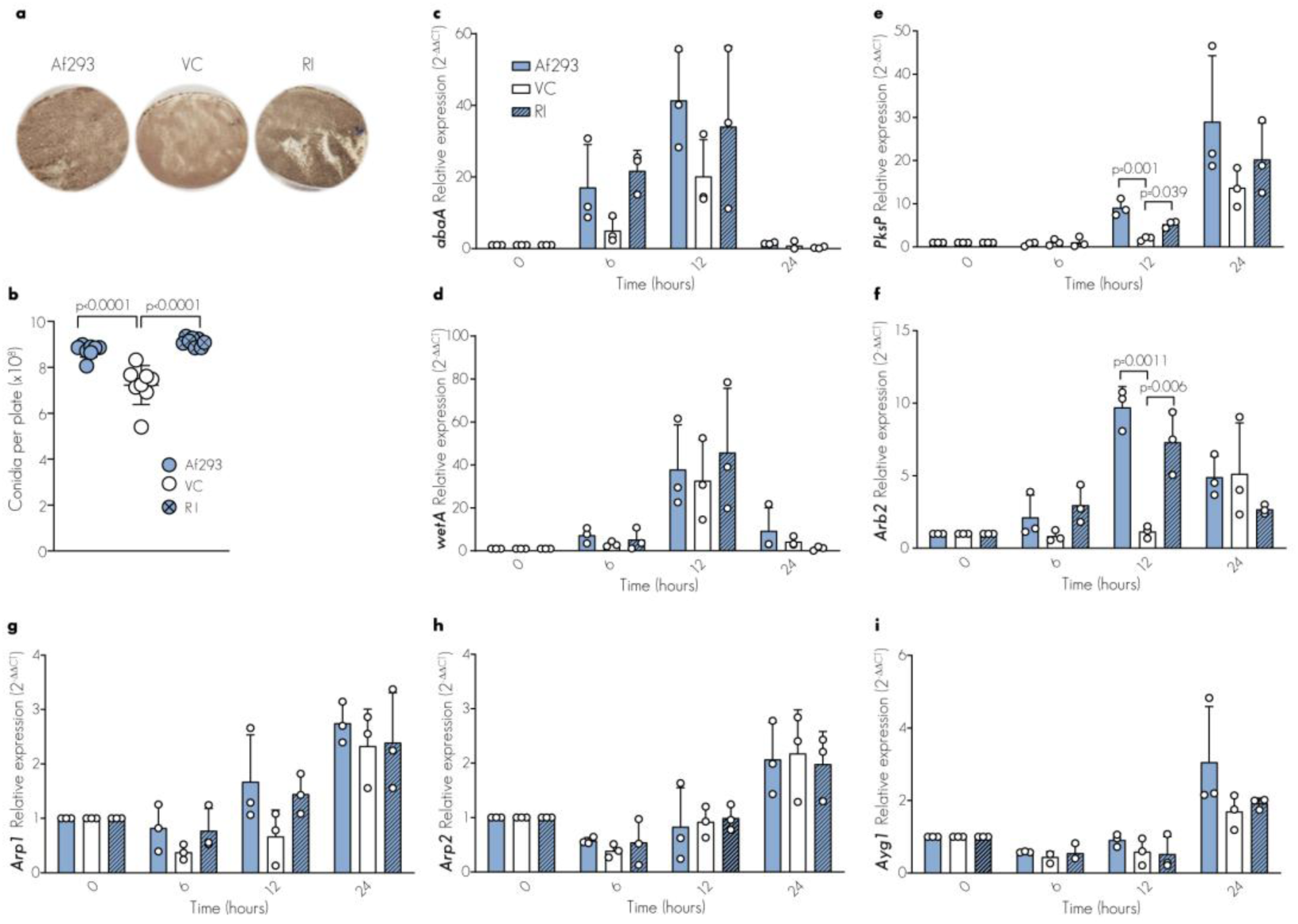
AfuPmV-1M regulates conidiogenesis. **a.** The AfuPmV-1M-infected strain (Af293) exhibits darker colonies compared to the ribavirin virus-cured strain. 2×10^5^ conidia (CFUs) were spread over GMM plates and grown for 7 days. **b**. Quantification of conidia number per plate on day 7. **C-i**. Real-time qPCR showing relative expression levels of (**c**) *abaA*, (**d**) *wetA*, (**e**) *PksP*, (**f**) *Arb2*, (**g**) *Arp1*, (**h**) *Arp2*, and (**i**) *Ayg1* during synchronized asexual developmental induction for *A. fumigatus Af293* and congenic strains. RNA was extracted at the indicated time points following the transfer of mycelium from the liquid-submerged synchronized culture to solid medium. Data were analyzed using the 2^−ΔΔCt^ method, with mycelium from the liquid-submerged synchronized culture as the control sample and normalized against 18S expression (reference gene). Conidial suspensions were adjusted based on viability staining to account for differences in conidial viability prior to each experiment. The data are derived from a minimum of three biologically independent experiments.

**Figure S4.**
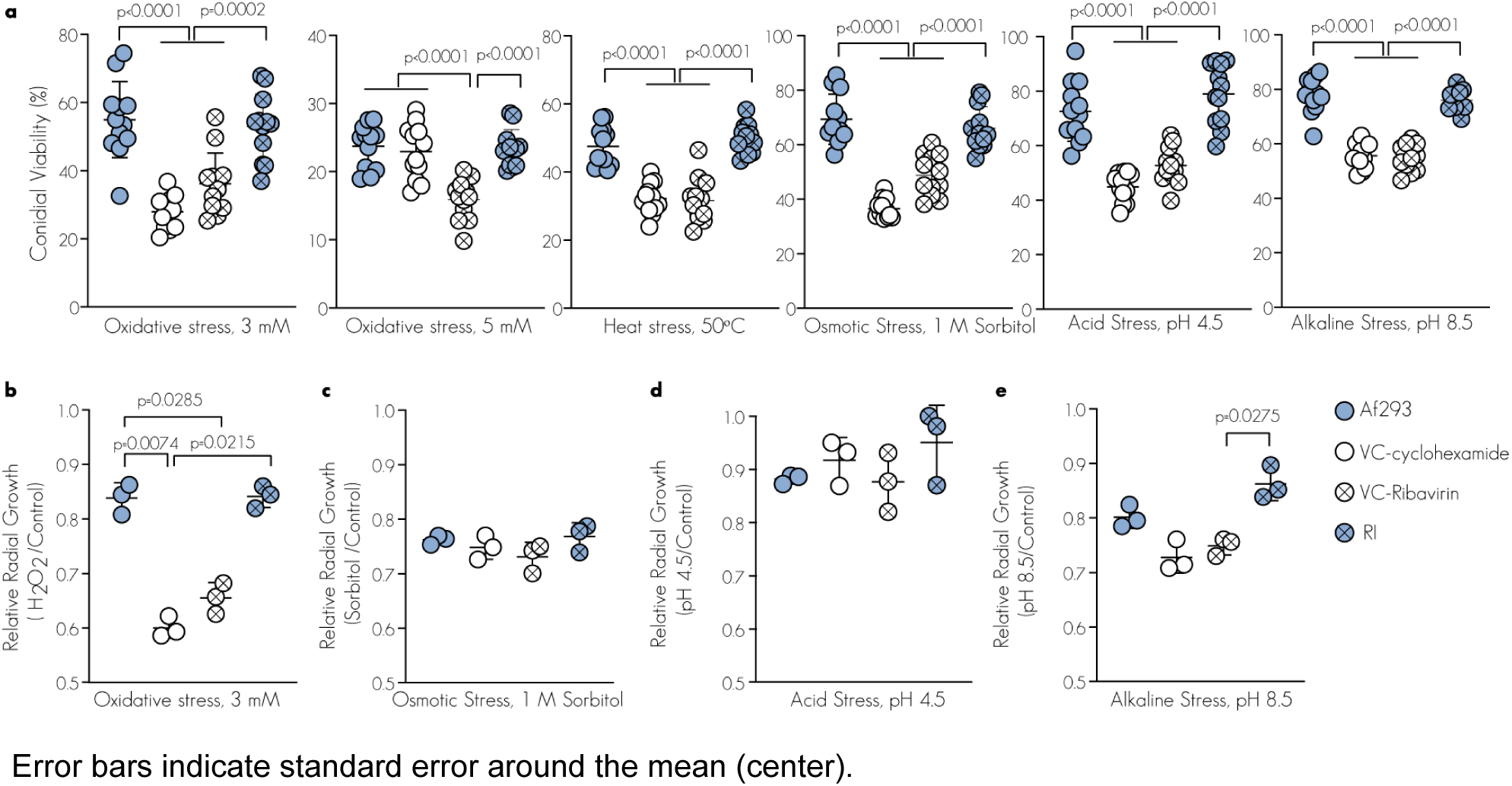
conidial viability. **a.** Conidial viability under stress conditions. Resting 2×10^5^ *Af293*, cycloheximide virus-cured, ribavirin virus-cued, and virus-re-infected (RI) conidia were exposed to 3 or 5 mM H2O2, 50°C, 1 M sorbitol, pH 4.5 or pH 8.5 for additional 4 h and analyzed for CFUs. **b-e**. Relative radial growth of 10^4^ conidia after five days incubation on solid GMM with 3 mM H_2_O_2_ (b), 1 M sorbitol (c), at pH=4.5 (d), or at pH=8.5 (e). Conidial suspensions were adjusted based on viability staining to account for differences in conidial viability prior to each experiment. The data are derived from a minimum of three biologically independent experiments. Statistical analysis: one-way ANOVA with Tukey’s multiple comparison test. Error bars indicate standard error around the mean (center). **Dataset 1.** Total spectra of *A. fumigatus* virus-infected and virus-cured strains using *Aspergillus fumigatus Af293* database.

**Figure S5.**
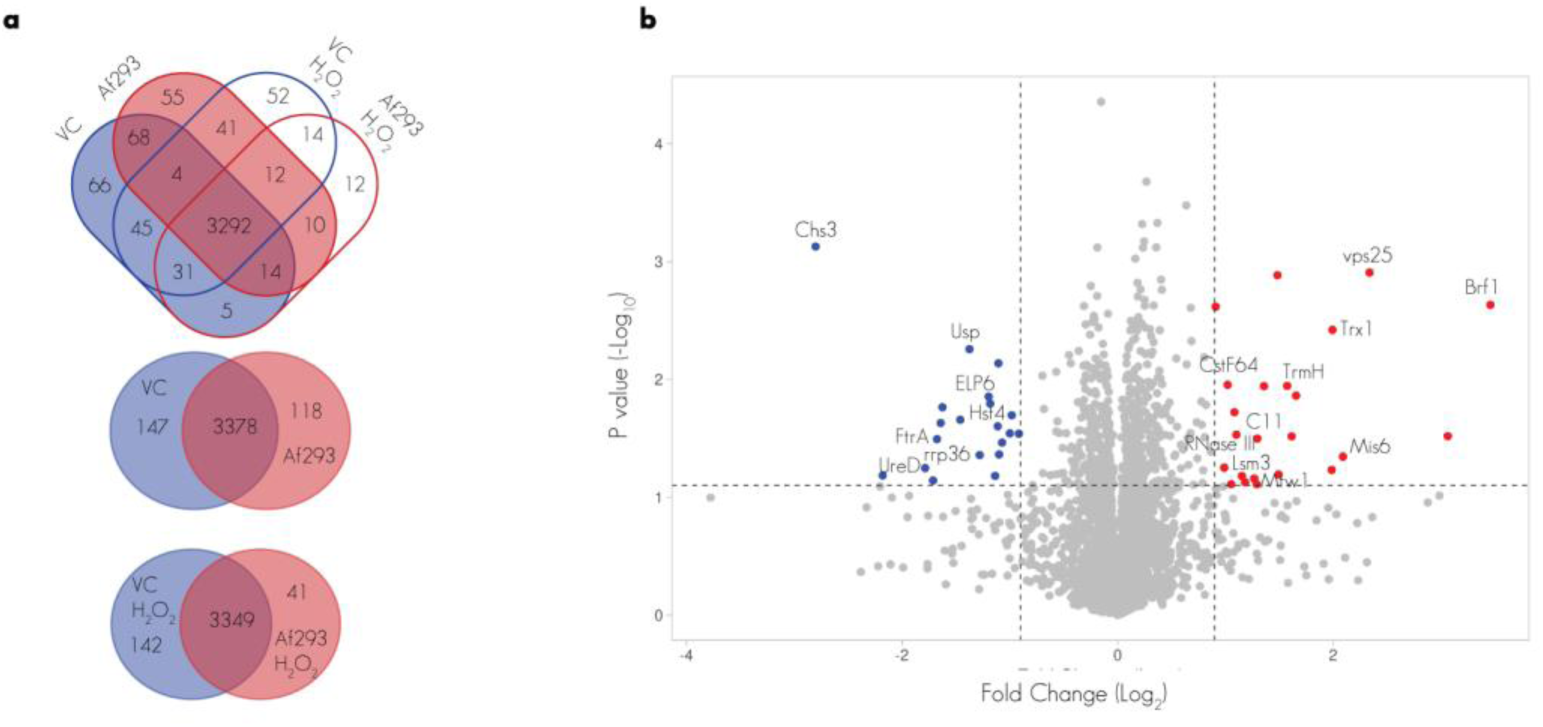
The proteome of AfuPmV-1M-infected *A. fumigatus* is altered. **a**. The protein content of *A. fumigatus* Af293 and cycloheximide virus-cured (VC) was characterized using liquid chromatography mass spectrometry (LC-MS/MS) under physiological and oxidative stress (8 mM) conditions. The distribution of the identified proteins by their presence/absence. Only proteins that appeared in 3 out of 3 triplicates (n = 3) from each group were included in the final list. **b**. The volcano plot shows the −log10 FDR as a function of log2 FC when comparing the Af293 virus-infected strain with the virus-cured strain (VC) grown under control conditions.

**Figure S6.**
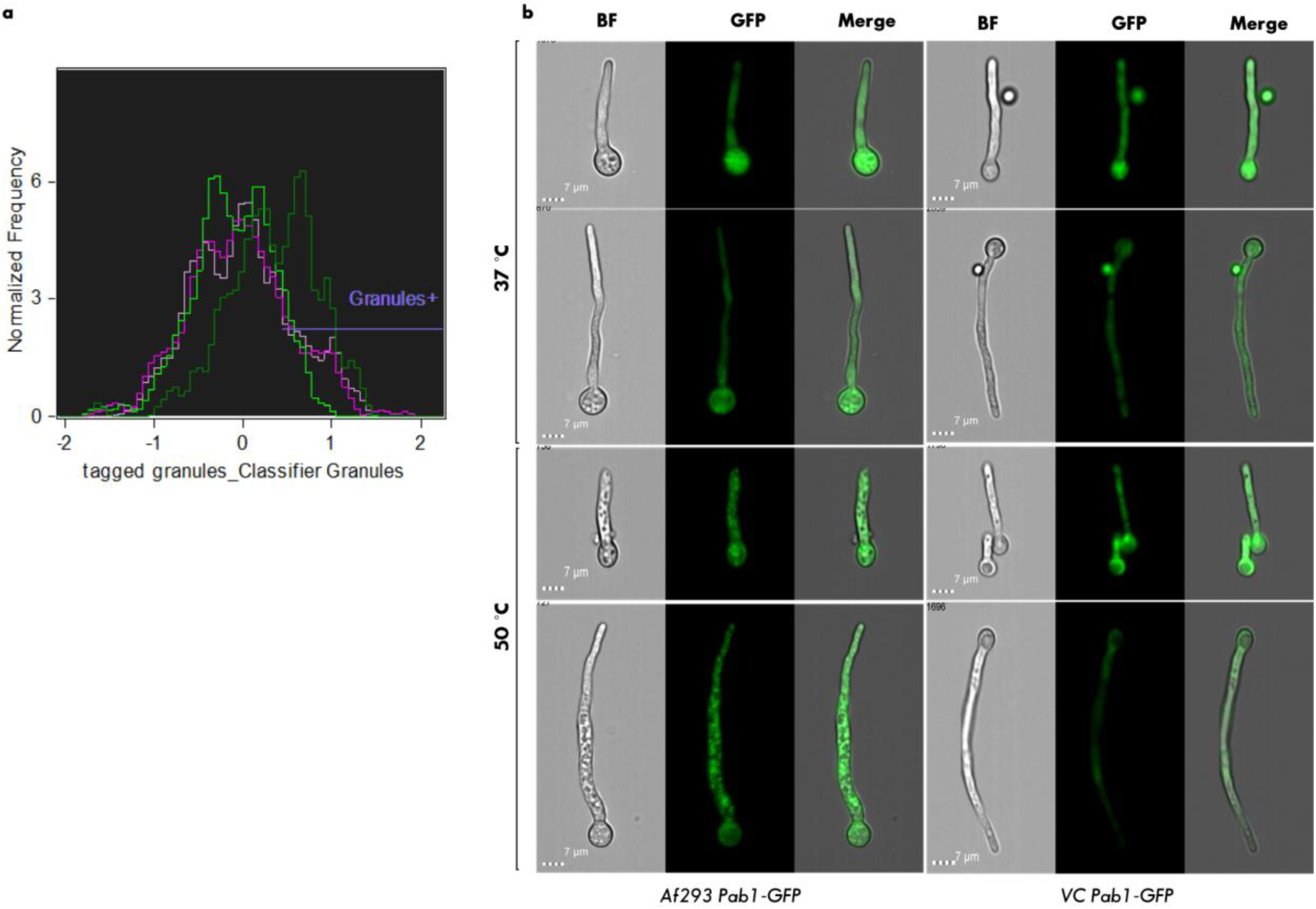
Targeting Afupmv-1M interfere with stress granule formation. Pab1-GFP germlings (7 h) were exposed to 50°C for 20 min and automated quantification of stress granule formation on gated cells was performed using the ImageStream technology. **a.** Representative histograms showing granularity **b.** Representative ImageStream images of PAB1-GFP expressing cells under control conditions (37°C) or upon heat shock stress (50°C, 20 min).

**Figure S7.**
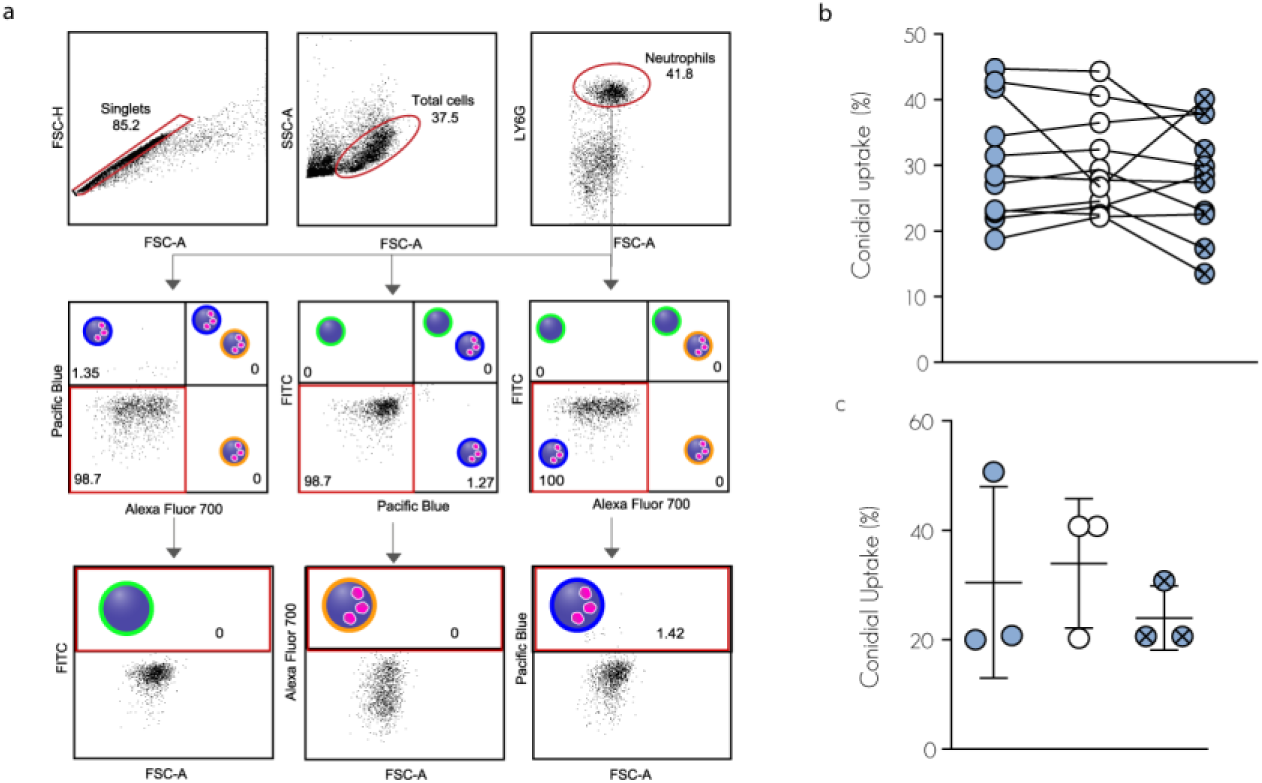
*A. fumigatus* mixed infection model. **a.** The gating strategy for isolating neutrophils involved utilizing lung samples from an unexposed animal to establish the criteria for identifying the positive cell populations. **b.** Uptake of virus-infected and virus-cured conidia by sorted murine neutrophil. The lines indicate paired data from a single mouse. The data are derived from three biologically independent experiments, each conducted with a cohort of four mice. **c**. Uptake of swollen conidia by environmental phagocytes *Acanthamoeba castellanii*. The data are derived from four biologically independent experiments, each performed in triplicate. Error bars indicate standard error around the mean (center).

**Figure S8.**
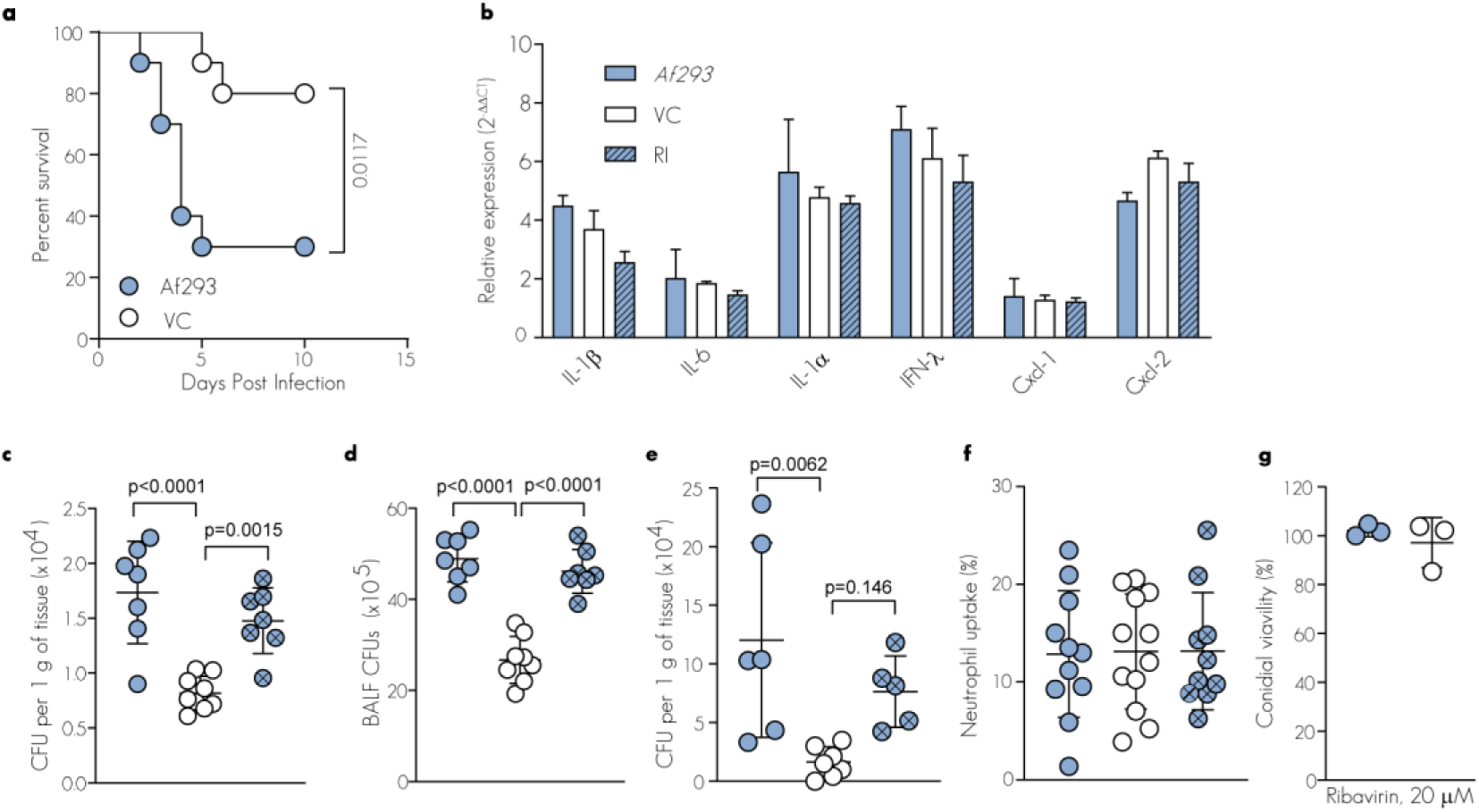
AfuPmV-1 regulates *A. fumigatus* virulence. **a.** Survival of C57BL/6JOlaHsd mice challenged with 6×10^7^ *Af293* (n=10) or virus-cured by ribavirin (VC, n=10) conidia. P values were calculated using log-rank Mantel–Cox test. **b.** Relative mRNA expression levels of proinflammatory mediators were assessed via RT-qPCR using total RNA extracted from human-derived macrophages at 6 hpi with the indicated strains. The data are pooled from two independent experimental replicates, normalized to actin expression, and the presented values are relative to the expression levels in naïve macrophages. **c**-**e**. C57BL/6J (**c-d**) and ICR mice (**e**) were challenged intranasally with 3×10^7^ Af293, virus-cured by ribavirin, or virus re-infected conidia. C57BL/6J lung (**c**) and BALF (**d**) CFUs at 24 hpi. **e**. ICR lung CFUs at 72 hpi. Samples are pooled from 2 independent experimental repetitions. **f**. Conidial uptake by C57BL/6J neutrophils at 24 hpi**. g.** Swollen virus-infected (Af293) or cycloheximide virus-cured conidia (4 h) were exposed to 20 µM ribavirin for an additional 4 h and analyzed for CFUs.

---

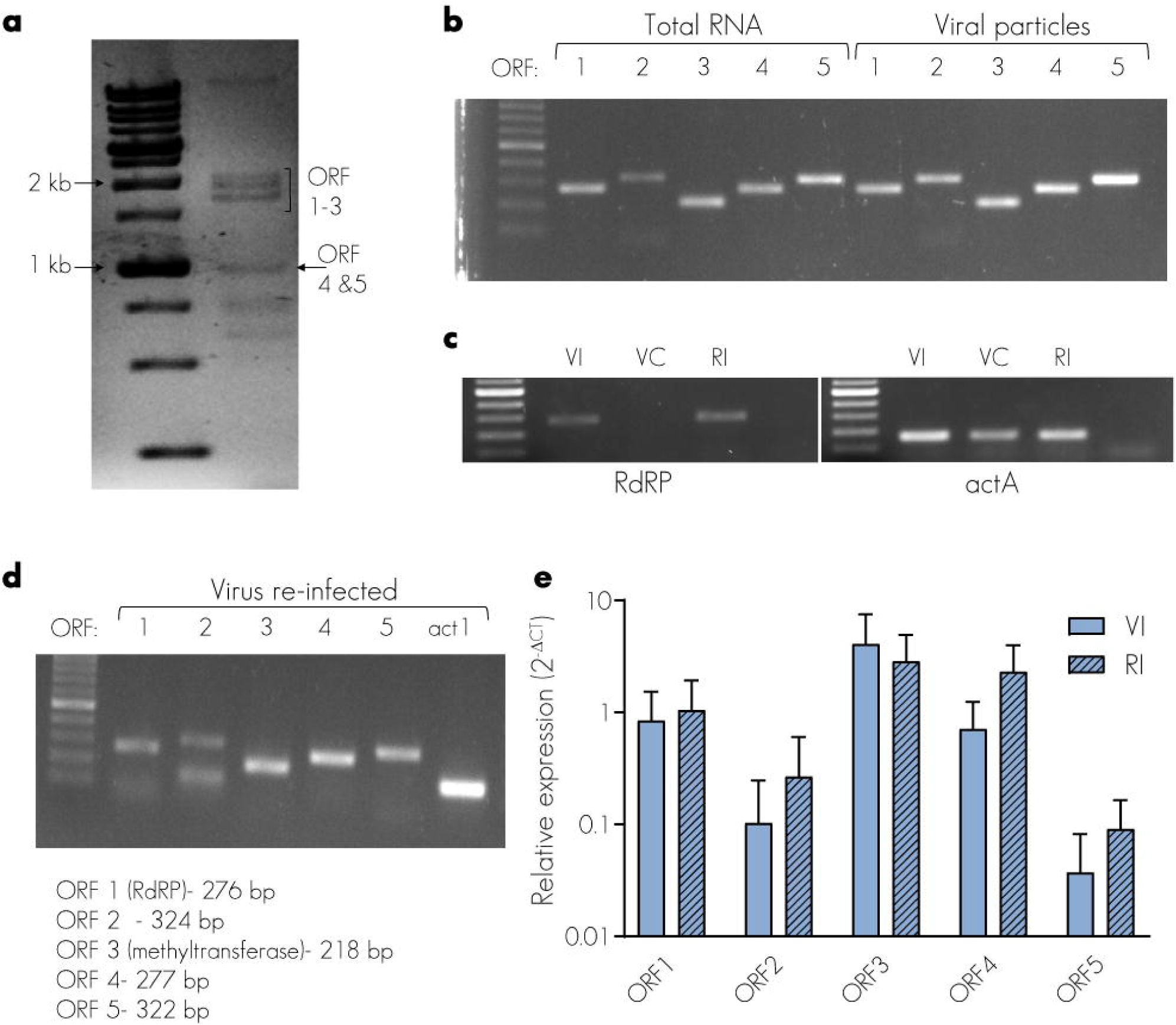

